# A Grass-Specific Structural Feature of Myosin VIII Regulates Protoxylem Development and Hydraulic Conductance in Sorghum

**DOI:** 10.64898/2026.02.27.708597

**Authors:** Zhiyuan Liu, Ran Tian, Leonidas D’Agostino, Zhanguo Xin, Junping Chen, Gunvant Baliram Patil, Yinping Jiao

## Abstract

Sorghum (*Sorghum bicolor*), a C₄ grass adapted to hot semi-arid environments, depends on reinforced xylem vessels to maintain hydraulic conductance under high evaporative demand. During protoxylem differentiation, coordinated microtubule and actin dynamics guide secondary cell wall (SCW) deposition; however, whether actin-based motors directly regulate vascular architecture and hydraulic performance has remained unknown. Here, we identify HEAT-SENSITIVE 1 (HS1), a previously uncharacterized myosin VIII, as a central regulator of protoxylem integrity and water transport in sorghum. The *hs1* mutant exhibited severe leaf scorching under field conditions. The *hs1* mutant displayed severe leaf scorching under field conditions, accompanied by pronounced protoxylem defects under controlled environments, including vessel collapse, reduced lumen area, and attenuated lignified SCWs. These structural abnormalities compromised longitudinal hydraulic conductance, diminished whole-plant water use, and rendered expanding leaves unable to meet transpirational demand, resulting in transient water deficit and thermal injury. *HS1* encoded a grass-specific myosin VIII with a grass-specific N-terminal extension exhibiting high intrinsic disorder and lineage-specific substitutions in the motor domain. Molecular and single-cell transcriptome analyses positioned *HS1* within differentiating protoxylem cells of developing leaves, revealing pronounced temporal and cell-type specificity. Furthermore bulk transcriptome profiling and quantitative lignin measurements indicated that *HS1* promoted lignin-associated SCW biosynthesis during protoxylem differentiation, functionally linking actin-based motor activity to wall reinforcement. *HS1* encodes a grass-specific myosin VIII distinguished by an extended intrinsically disordered N-terminal region and lineage-specific substitutions within the motor domain, suggesting evolutionary specialization. Consistant with core developmental role, reduced nucleotide diversity at the *HS1* locus across diverse sorghum accessions further supported strong evolutionary constraint. Together, these findings established HS1 as the first actin-based motor protein shown to control xylem architecture and hydraulic function in plants.

## INTRODUCTION

Sorghum is a diploid C₄ grass of the *Poaceae* family, the fifth most produced cereal crop worldwide (Fontanet-Manzaneque et al., 2025; Nigam et al., 2025; Zhang et al., 2025). Among the major grass crops, such as wheat, rice, maize, and barley, sorghum exhibits outstanding heat and drought tolerance, making it a critically important crop for agricultural production in arid and semi-arid regions (House, 1985; Parent and Tardieu, 2012). Understanding the mechanistic basis of this resilience has become crucial under increasingly adverse environmental conditions in crop production.

The highly efficient xylem hydraulic system is a kye contributor to sorghum’s tolerance to heat and drought stress. This efficiency is largely determined by xylem anatomical features such as vessel diameter and secondary cell wall (SCW) thickness which support water transport efficiency, limits cavitation, and sustains transpiration (Drobnitch et al., 2021; Lehrer and Hawkins, 2023; Torres-Ruiz et al., 2024). Hydraulic efficiency ultimately depends on the coordinated development of xylem cell types with lignified SCW, which provide mechanical strength, structural integrity and prevent conduit collapse (Rogers and Campbell, 2004; Hwang et al., 2016). Among these, protoxylem represents the earliest differentiated xylem tissue and play a critical role in initiating water transport during early organ development, particularly in the developing leaf blades (Goodwin, 1942; Sharman, 1942). Xylem development is orchestrated through multiple, tightly coordinated regulatory layers that operate across distinct developmental stages. Hormonal signaling and ubiquitin-mediated pathways predominantly regulate early vascular cell fate decisions and differentiation competence (Köllmer et al., 2014; Fabregas et al., 2015; Zhang et al., 2016; Smetana et al., 2019; Phookaew et al., 2024).

As differentiation progresses, xylem maturation is governed by complex and highly integrated regulatory networks. These include hierarchical transcriptional programs driven by NAC master regulators such as VND7 and SND1 (Kubo et al., 2005; Yamaguchi and Demura, 2010) and their downstream MYB factors including MYB46 (Zhong et al., 2007; Xiao et al., 2021), coordinated activation of enzymes responsible for SCW biosynthesis (Turner and Somerville, 1997; Taylor et al., 2004; Wu et al., 2009), cytoskeleton-dependent spatial control of SCW deposition (Oda and Fukuda, 2013; Higa et al., 2024), and tightly regulated hydrolase activities that execute programmed cell death to generate functional conduits (Groover and Jones, 1999; Funk et al., 2002)

Cytoskeletal dynamics constitute a critical but understudied regulatory layer in xylem differentiation, linking transcriptional programs to the spatial patterning of SCWs. Cytoskeletal dynamics during xylem differentiation are driven by motor proteins, including kinesins and myosin (Nebenführ and Dixit, 2018; Ryan and Nebenführ, 2018). To date, only two kinesins have been functionally correlated in xylem development: Kinesin-13A, which regulates metaxylem pit size, and FRAGILE FIBER1 (FRA1), which controls protoxylem SCW thickness through microtubule organization (Oda and Fukuda, 2013; Zhu et al., 2015). Both genes modulate microtubule stability and organization during SCW formation in *Arabidopsis*. Despite these roles in xylem patterning, direct evidence linking kinesin activity to hydraulic function is lacking, and no myosin has yet been functionally characterized for xylem development or water transport.

Myosins are evolutionarily conserved actin-based motor proteins that drive intracellular transport and spatial organization of cellular components (Madison and Nebenführ, 2013). In the plants, the myosin superfamily comprises two major classes, myosin XI and myosin VIII, which differ in abundance, localization and biological function (Peremyslov et al., 2011). Myosin XI proteins have been shown to contribute to stem proprioception in *Arabidopsis*, protein body formation during maize endosperm development, and powdery mildew resistance in barley (Wang et al., 2012; Okamoto et al., 2015; Acevedo-Garcia et al., 2022). To date, functional characterization of myosin VIII genes remains limited. Studies in moss have linked myosin VIII to cell division and developmental patterning (Wu et al., 2011; Wu and Bezanilla, 2014), while in *Arabidopsis* several myosin VIII isoforms facilitate *Agrobacterium*-mediated transformation (Liu et al., 2023). *Arabidopsis thaliana* MYOSIN 1 (ATM1; AT3G19960) is the only myosin VIII member shown to regulate a developmental function, controlling root meristem cell proliferation via auxin and sugar signaling pathways (Olatunji et al., 2023). To date, no additional myosin VIII gene has been causally linked to a defined plant trait at the single-gene level. Here, we identify a grass-specific class VIII myosin as a key regulator of protoxylem integrity and water transport in sorghum, a crop renowned for its resilience to heat and drought. This study reveals an evolutionarily specialized function of myosin VIII that is absent from *Arabidopsis*, highlighting how cytoskeletal regulation has diversified in grasses to support distinct vascular and hydraulic demands. By linking actin-based motor activity to secondary cell wall integrity and xylem performance, our findings extend cytoskeletal paradigms from dicot models to grass vascular biology and uncover a previously unrecognized mechanism contributing to hydraulic robustness and environmental adaptation.

## RESULTS

### Sorghum heat-sensitive mutant *hs1*

A heat-sensitive sorghum mutant, *hs1,* was isolated from an EMS-induced mutant population generated in the genetic background of reference line BTx623 (Xin et al., 2008; Jiao et al., 2016; Jiao et al., 2024). The mutant was initially designated *heat sensitive 1* based on the consistent appearance of leaf scorching of developing leaves after an exposure to a moderate heat stress event (>34°C) under well-watered field conditions at or after 5-leaf stages and throughout the vegetative developmental stages (Fig. 1A; Supplementary Figs. S1, A and B). The scorching phenotype was initiated at the top part of the developing leaf and moved downward to the mid to lower sections of the leaf. The severity of leaf phenotypes and the number of leaves being affected were correlated to the intensity and duration of a particular heat wave (Supplementary Fig. S1C). Therefore, multiple leaves showed a leaf-scorching phenotype after multiple heat stress events during the growing season (Fig. 1A; Supplementary Figs. S1, A to C). As a result of heat-induced permanent leaf damage, *hs1* showed significant reductions in leaf length and plant height compared to wild-type (WT) grown under the same environments (p < 0.05; Figs. 1, B and C; Supplementary Figs. S1, D to G). In addition, the reproductive tissues of *hs1* are also sensitive to heat stress. Heat stress at early stages of panicle development reduced panicle length, at the half bloom stage caused panicle blast (Fig. 1D), and at the pollination stage significantly decreased seed numbers (p < 0.01; Figs. 1, E and F).

**Figure 1.**
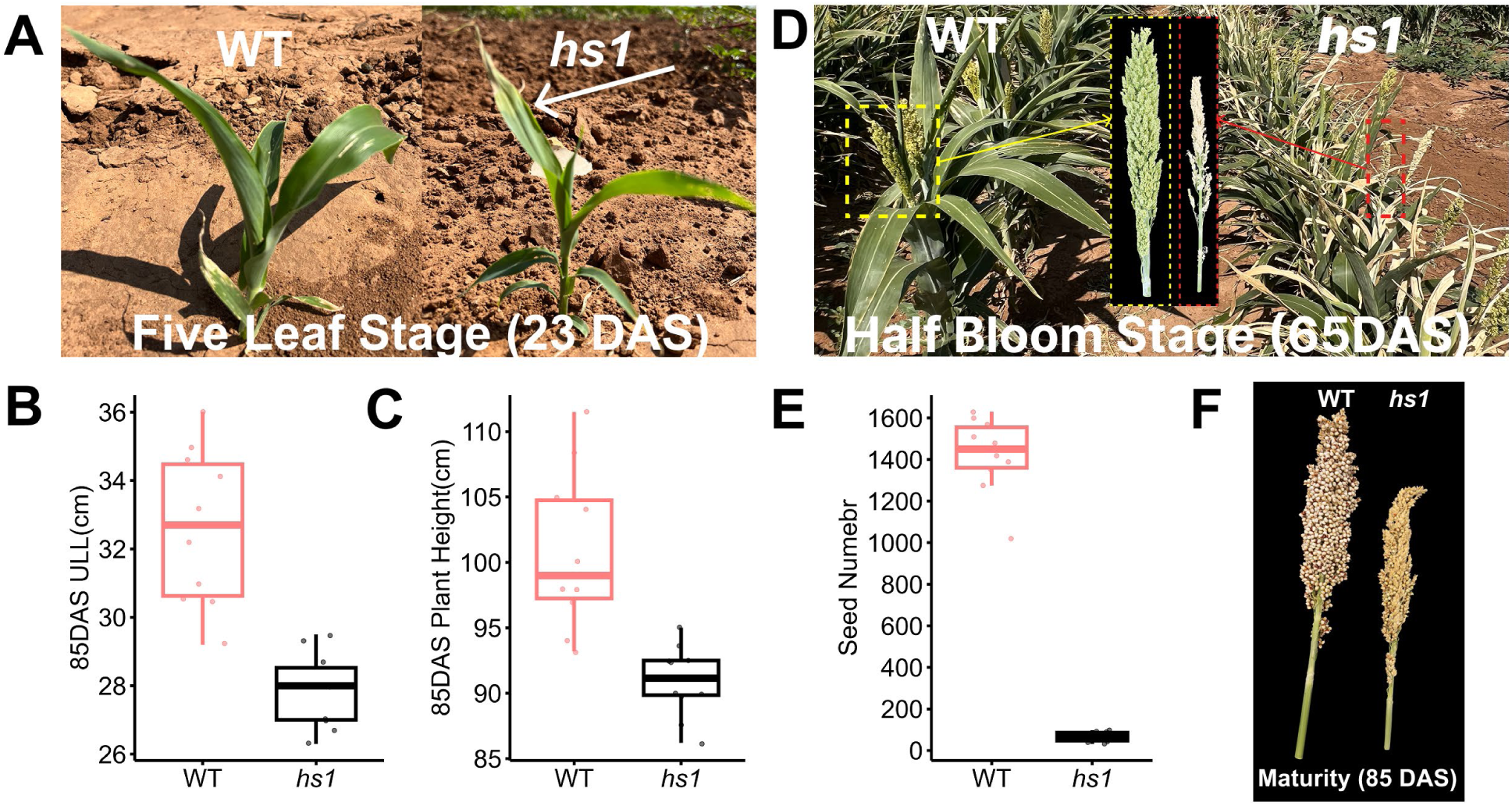
Phenotype of heat-sensitive mutant *hs1*. **A)** Leaf phenotype of wild type (WT, BTx623) and *hs1* grown in a field (Lubbock, TX) at the five-leaf stage (23 days after sowing, DAS). The white arrow indicates a scorched developing leaf. **B, C)** Quantification of upper leaf length (ULL) (B) and plant height (C) at 85 DAS. **D)** Panicle phenotype of WT and *hs1* at the half bloom stage; middle panel shows a close-up view. **E, F)** Quantification of the seed number at the physiological maturity. For statistical analyses, boxes indicate the first and third quartiles, and horizontal lines indicate the median (n = 10). **P < 0.01 (two-tailed Student’s t-test).

Another phenotype in the field related to *hs1* was leaf roll of developing leaves in the afternoon under low relative humidity (RH; 29–33%). The rolled leaf partially recovered the next morning but reappeared in the hot afternoons (Supplementary Fig. S2A). Overtime, the rolled leaf became bleached grey in color, leading to permanent leaf damage after repeated daily heat exposure (Supplementary Fig. S2A). Notably, similar leaf roll and loss of leaf greenness were also observed in developing *hs1* leaves during afternoon under naturally elevated light, although plants were maintained under optimal growth conditions for sorghum (defined as 26°C–30 °C with natural light ranging from 430 to 1162 μmol m⁻² s⁻¹ photosynthetic photon flux density (PPFD); Supplementary Fig. S1C).

The heat sensitive phenotypes of *hs1* observed in hot and dry environments were not observed in the winter nursery field in Guayanilla, Puerto Rico (18.0373°N, 66.7963°W, elevation 49 m), a high RH environment with no heat stress (20°C -28°C). The contrasting phenotypes between WT and *hs1*, as well as the differential performance of *hs1* across environments, suggested that impaired evaporative cooling and transpiration capacity contributed to temperature- and environment-dependent tissue damage and growth defects throughout development.

### Collapsed protoxylem vessels in developing leaves of *hs1* disrupted water transport

To determine the developmental defects underlying the scorching leaf phenotype, we examined root traits, stomatal characteristics, and protoxylem structure, which are key components of water uptake and transpiration, under optimal greenhouse conditions.

At the growing point differentiation (GPD) stage, root surface area and root sap (defined as root xylem sap exudate) did not differ significantly between WT and *hs1* (P > 0.05; Supplementary Figs. S3, A to D). Similarly, no differences were observed in stomatal size (measured as guard cell length) and density in developing leaves between the two genotypes (P > 0.05; Supplementary Figs. S4, A to D), indicating that neither root water uptake capacity nor stomatal traits account for the observed phenotype. In contrast, safranin uptake assay using cut stems revealed clear differences in water transport between WT and *hs1*. The volume of solutions absorbed by WT was significantly greater than that by *hs1* under naturally elevated light conditions (P < 0.05; Supplementary Figs. S5, A and B). Although mature leaves of both genotypes exhibited similar staining after 2 h, developing *hs1* leaves showed markedly reduced or absent dye accumulation (Supplementary Figs. S4, C to E).

When developing ninth leaves were detached from the whole plant for analysis, dye uptake and midrib transport distance were reduced by 23.1% and 27.3%, respectively, in *hs1* relative to WT (P < 0.05; Figs. 2, A to C). In *hs1*, staining stopped at the upper midrib, whereas WT leaves showed continuous, more intense staining along the veins (Fig. 2D). Together, these results demonstrate that the leaf-scorching phenotype of *hs1* arises from impaired internal water transport and distribution in developing leaves, rather than defects in water uptake or stomatal regulation.

**Figure 2.**
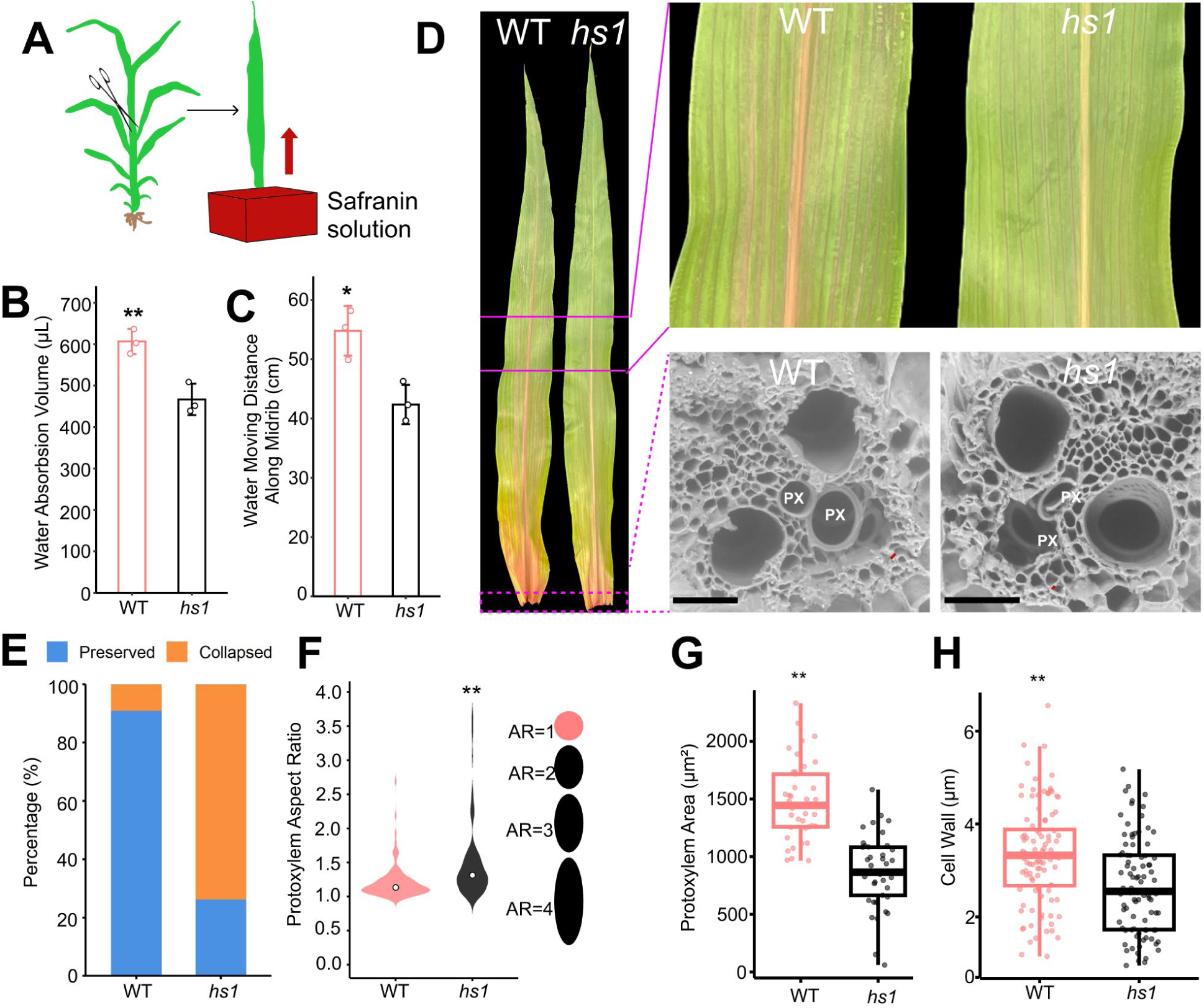
Impaired water transport in developing leaves of *hs1*. **A)** Schematic of the safranin uptake assay used to assess xylem-mediate water transport. **B, C)** Quantification of safranin dye uptake (B) and midrib transport distance (C) in WT and *hs1*. Values represent mean ± SD (n = three biological replicates per genotype). **D)** Representative developing leaves of WT and *hs1* following the safranin assay. Upper right, enlarged view of leaves. Lower right, scanning electron microscopy (SEM) images of midrib transverse sections from the leaf base. PX, protoxylem. Bars = 50 µm. **E to H)** Quantification of protoxylem (PX) integrity in midrib transverse sections from the leaf base of developing WT and *hs1* plants, including frequency of collapsed versus intact vessels (E), aspect ratio (F), inner diameter (G), and secondary cell wall thickness (H). Data were obtained from three biological replicates per genotype. Individual PX vessels were pooled for analysis (n ≥ 42 for E; n ≥ 93 for F; n = 41 for G; n ≥ 90 for H). Boxes represent the first and third quartiles, and horizontal lines indicate medians. Statistical significance was determined using two-tailed Student’s t-tests (*P < 0.05; **P < 0.01).

As protoxylem mediates water transport in developing leaves, particularly in the elongating basal region (Goodwin, 1942), we examined midrib protoxylem structure at the developing leaf base. In WT, protoxylem vessels were intact and nearly circular, with secondary cell walls (SCW) confined to the vessel boundary (Fig. 2D). By contrast, *hs1* protoxylem showed a continuum of structural defects, including mild elliptical deformation, spindle-shaped distortion and SCW extrusion. In severe cases, disintegrated protoxylem remnants accumulated within the vessel cavity (Fig. 2D; Supplementary Figs. S6, A to D).

Quantitative analysis confirmed extensive structural defects in *hs1*. Approximately 73.8% of protoxylem vessels were collapsed in *hs1*, compared with 11.11% in WT (Fig. 2E). *hs1* vessels had significantly higher aspect ratios (P < 0.01; Fig. 2F), smaller lumen areas (870 ± 310 μm² vs. 1479 ± 332 μm² in WT; P < 0.01; Fig. 2G), and thinner secondary walls (2.66 ± 1.02 µm vs. 3.34 ± 1.05 µm in WT; P < 0.01; Fig. 2H).

To determine whether this vascular defect persists across developmental stages under optimal growth conditions, we next examined successive developing leaves. Leaf scorching was consistently observed in *hs1* across successive developmental stages under optimal growth conditions, including the sixth leaf at the five-leaf stage, the seventh through eleventh leaves at the GPD stage, and the flag leaf at the flag-leaf stage (Supplementary Fig. S7A). Detailed observation of these developing leaves (excluding the ninth, which was analyzed previously) revealed the presence of collapsed protoxylem at each stage (Supplementary Figs. S7A). Quantification revealed that the ratios of collapsed xylem in *hs1* ranged from 56-74%, which were higher than those in the WT (Supplementary Figs. S7B). Collectively, these defects compromise protoxylem integrity and provide a structural explanation for reduced water transport in developing *hs1* leaves.

### Physiological consequences of protoxylem collapse

We next evaluated the physiological consequences of protoxylem disruption of leaf water relations in the greenhouse under optimal conditions. During the morning, developing leaves of WT and *hs1* were visually indistinguishable and showed comparable H₂O₂ accumulation (Figs. 3, A and B). However, despite the absence of visible stress, *hs1* leaves exhibited lower stomatal conductance (gₛ) and transpiration rate (E) (P < 0.01; Supplementary Figs. S8, A and B), while relative water content, chlorophyll concentration, and electrolyte leakage remained indistinguishable from WT (P > 0.05; Figs. 3, C to E). These results indicate reduced hydraulic efficiency in *hs1* prior to visible symptoms.

**Figure 3.**
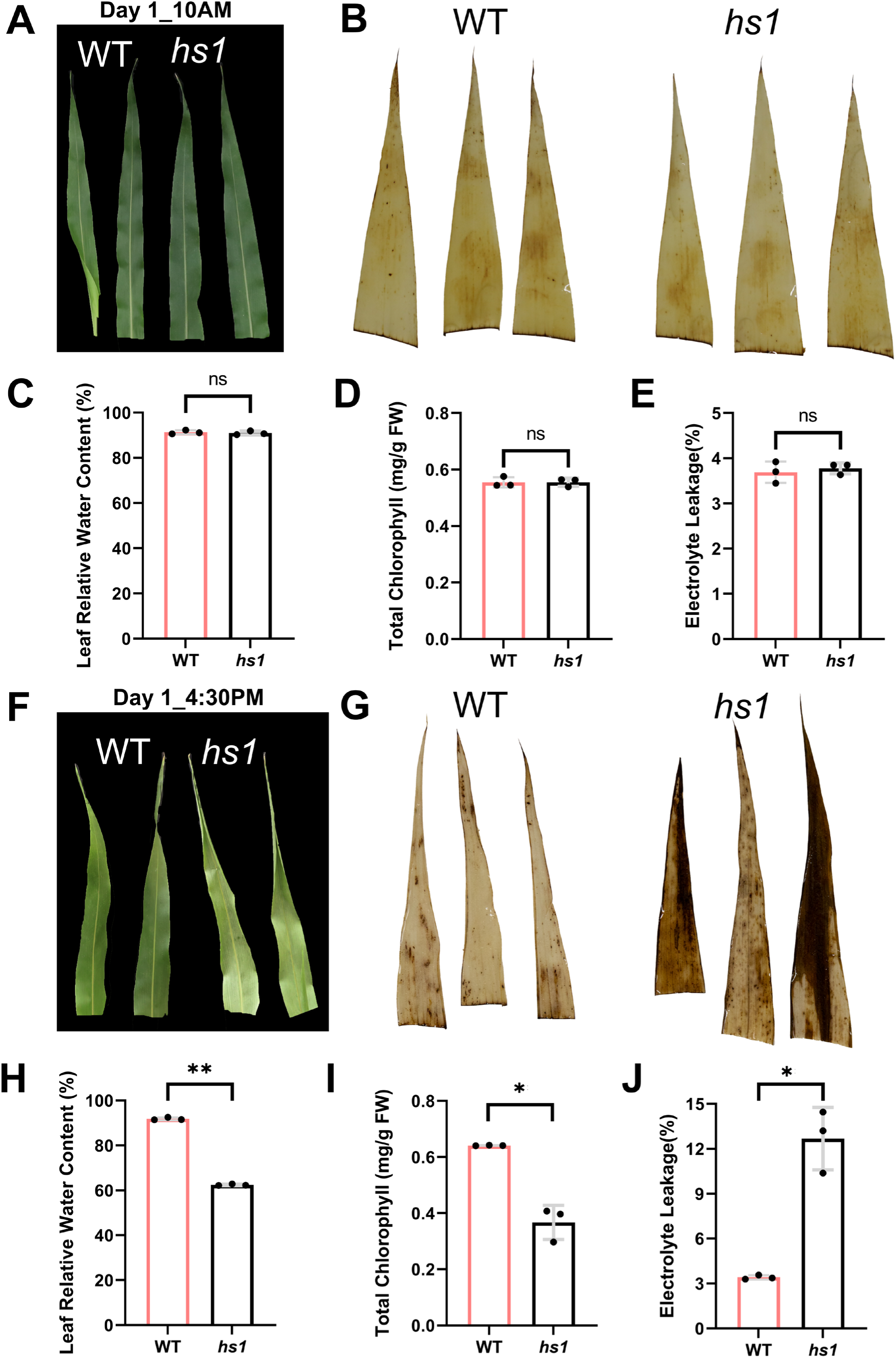
Morphology and physiology characterization of developing leaves in WT and *hs1*. **A, B)** Phenotypic comparisons of developing leaf morphology (A) and 3,3′-diaminobenzidine (DAB) staining (B) in WT and *hs1* plants at 10 AM. **C to E)** Statistical comparisons of leaf relative water content (C), total chlorophyll content (D), and electrolyte leakage (EL) (E) in WT and *hs1* at 10AM. **F, G)** Phenotypic comparisons of developing leaf morphology (F) and DAB staining (G) in WT and *hs1* plants at 4:30 PM. **H to J)** Statistical comparisons of leaf relative water content (H), total chlorophyll content (I), and EL (J) in WT and *hs1* plants at 4:30 PM. For all statistical analyses, values are shown as the mean ± standard deviation from three biological replicates. **P < 0.01, *P < 0.05, ns, not significant P>0.05 (two-tailed Student’s t-test).

As evaporative demand increased in the afternoon (under higher light and temperature), WT leaves significantly increased gₛ and E (all P<0.01) to meet transpiration demand (Supplementary Figs. S9, A and B), whereas *hs1* leaves showed a significant decline (all P < 0.05; Supplementary Figs. S9, C and D), accompanied by visible scorching and stronger DAB staining in developing leaves (Figs. 3, F and G). Consistently, *hs1* leaves exhibited significantly lower gₛ and E, reduced relative water content and chlorophyll, and increased electrolyte leakage (all P < 0.01; Figs. 3, H to J; Supplementary Figs. S9, E and F). Approximately 73% of scorched *hs1* leaves recovered the chlorophyll content after two days, whereas WT leaves remained unaffected (Supplementary Figs. S9, A and B). At the whole-plant level, *hs1* plants exhibited significantly lower daily water uptake and reduced stature compared with WT (all P < 0.01; Supplementary Figs. S11, A to C). Detached leaf assays further showed slower initial water loss in *hs1* (Supplementary Figs. S11, D to F), consistent with impaired hydraulic conductance.

Together, these results demonstrated that collapse of protoxylem vessels in developing *hs1* leaves compromises vascular integrity and reduces hydraulic efficiency. This defect was detectable before visible symptoms and limited the ability of developing leaves to meet increased transpirational demand during afternoon periods of higher light and temperature. Consequently, *hs1* leaves experienced transient water deficit, oxidative stress, and leaf scorching, establishing protoxylem integrity as a critical determinant of water transport during leaf development.

### *HS1* encodes a myosin protein

To determine whether the causal mutation of the *hs1* is dominant or recessive, *hs1* was crossed with WT to generate an F₁ population. All 43 individuals exhibited a WT phenotype without any leaf damage in the field condition, indicating that the causal mutation is recessive (Supplementary Fig. S12A). In subsequent F₂ population, leaf scorching and WT phenotypes segregated at a ratio of 20:73, consistent with the expected 1:3 Mendelian ratio (χ² = 0.61, P = 0.44). Together, these segregation patterns demonstrated that the leaf scorching phenotype in *hs1* is governed by a single recessive mutation.

Next, bulked segregant analysis (BSA) of the F₂ population was performed with four biological replicates over two years to identify the causal mutation (Supplementary Fig. S12B). Two genes (*Sobic.002G340200* and *Sobic.002G339300*) on chromosome 2 were consistently identified in the four replicates. Screening alternative alleles of these two genes across our sequenced 1,256 mutants (Jiao et al., 2024) revealed that only *Sobic.002G339300,* carried four independent alleles producing a similar leaf scorching phenotype (Fig. 4A, Supplementary Fig. S12C). To further validate *Sobic.002G339300* as the causal gene, a complementation (cis–trans) test was conducted by crossing homozygous *hs1* with the mutant *25M2-1883*. All F₁ offspring displayed the leaf scorching phenotype (Fig. 4B), confirming that *Sobic.002G339300* is the causal gene, designated *HS1*.

**Figure 4.**
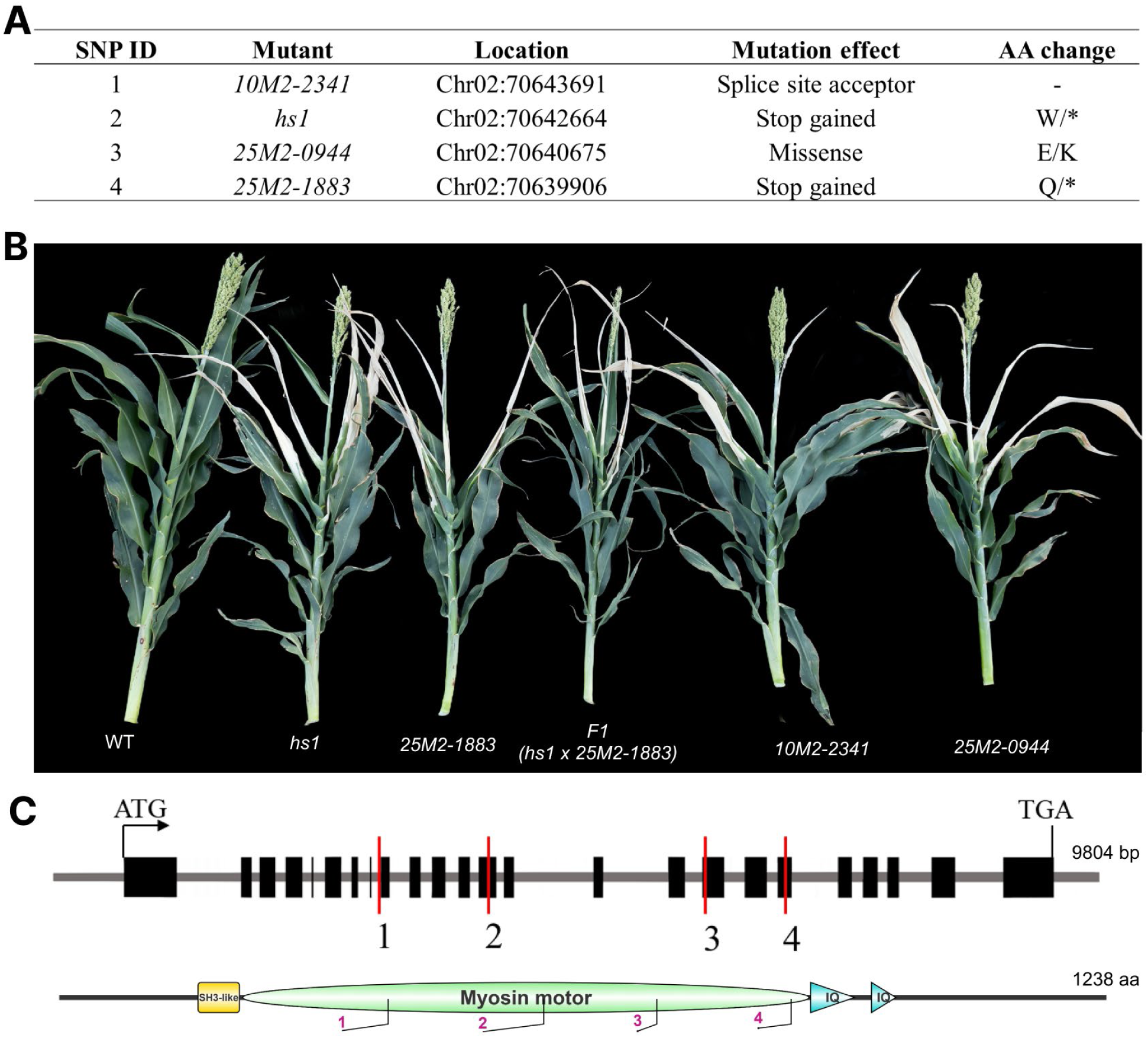
Identification of the causal gene for the *hs1* mutant. **A)** Four independent alleles of the candidate gene *Sobic.002G339300.* B) Complementation test to validate the causal gene. Representative phenotypes of WT, *hs1*, *25M2-1883*, the F1 progeny from the *hs1* × *25M2-1883* cross, and two additional independent mutant alleles (*10M2-2341* and *25M2-0944*) showing leaf scorching. **C)** Gene structure and protein domains of *Sobic.002G339300* (myosin protein). Positions of the four independent mutations shown in (A) are indicated.

The WT *HS1* gene encodes a myosin protein and consists of three major functional domains: src Homology-3 (SH3)-like, Motor domain, and isoleucine–glutamine (IQ) motif containing neck region (Fig. 4C). Structural modeling of the *hs1* mutant type protein revealed substantial disruption of protein architecture, including truncation of the tail region and pronounced disorder within the head and neck regions (Supplementary Figs. S13, A to C). The *hs1* allele introduced a premature stop codon at the 573^rd^ amino acid, and structural alignment with the WT protein revealed significant conformational deviation (TM-score = 0.34/0.73; RMSD = 1.22 Å). These variants in protein structures strongly suggest impaired *HS1* motor function in the mutant background.

### *HS1* encodes a myosin VIII with a grass-specific structural feature

Plant myosins are classified into two classes: myosin XI and VIII. The phylogenetic analyses of these classes across *Sorghum bicolor* (12 genes), *Zea mays* (8 genes), *Oryza sativa* (8 genes), and *Arabidopsis thaliana* (17 genes), classified *HS1* as a member of the myosin VIII class (Supplementary Fig. S14). We next expanded our phylogenetic survey across plants. No intact myosin VIII class proteins were detected in surveyed algal lineages (Supplementary Data Set 1), consistent with previous reports (Richards and Cavalier-Smith, 2005; Peremyslov et al., 2011; Mühlhausen and Kollmar, 2013; Sebe-Pedros et al., 2014; Nebenführ and Dixit, 2018). In contrast, myosin VIII was identified throughout land plants (Supplementary Data Set 1). We compiled 134 myosin VIII class proteins from 44 species spanning early-diverging land plants to angiosperms and constructed a phylogenetic tree using three *Arabidopsis* myosin XI class proteins as outgroup (Supplementary File 1).

The phylogenetic tree showed that myosin XI proteins branched outside the myosin VIII clade, defining the root of the myosin VIII lineage (Fig. 5A; Supplementary Fig. S15). Within myosin VIII class, lycophyte sequences occupied basal positions, followed by bryophytes, which formed a sister clade to angiosperms (Fig. 5A). Angiosperm myosin VIII proteins segregated into two major paralogous clades, designated VIII-A and VIII-B, reflecting an ancient duplication near the base of angiosperms (Peremyslov et al., 2011; Mühlhausen and Kollmar, 2013). Within each clade, basal angiosperm sequences preceded a clear divergence between monocot and eudicot lineages (Fig. 5A). Within each monocot lineage, grass sequences formed distinct subclades. *HS1* clustered within the grass-specific VIII-B clade, which contained 16 members from 11 grass species (Fig. 5A; Supplementary Fig. S15), compared with 17 members from 12 species in the grass VIII-A clade. These results indicate that *HS1* is a grass-specific myosin VIII protein.

**Figure 5.**
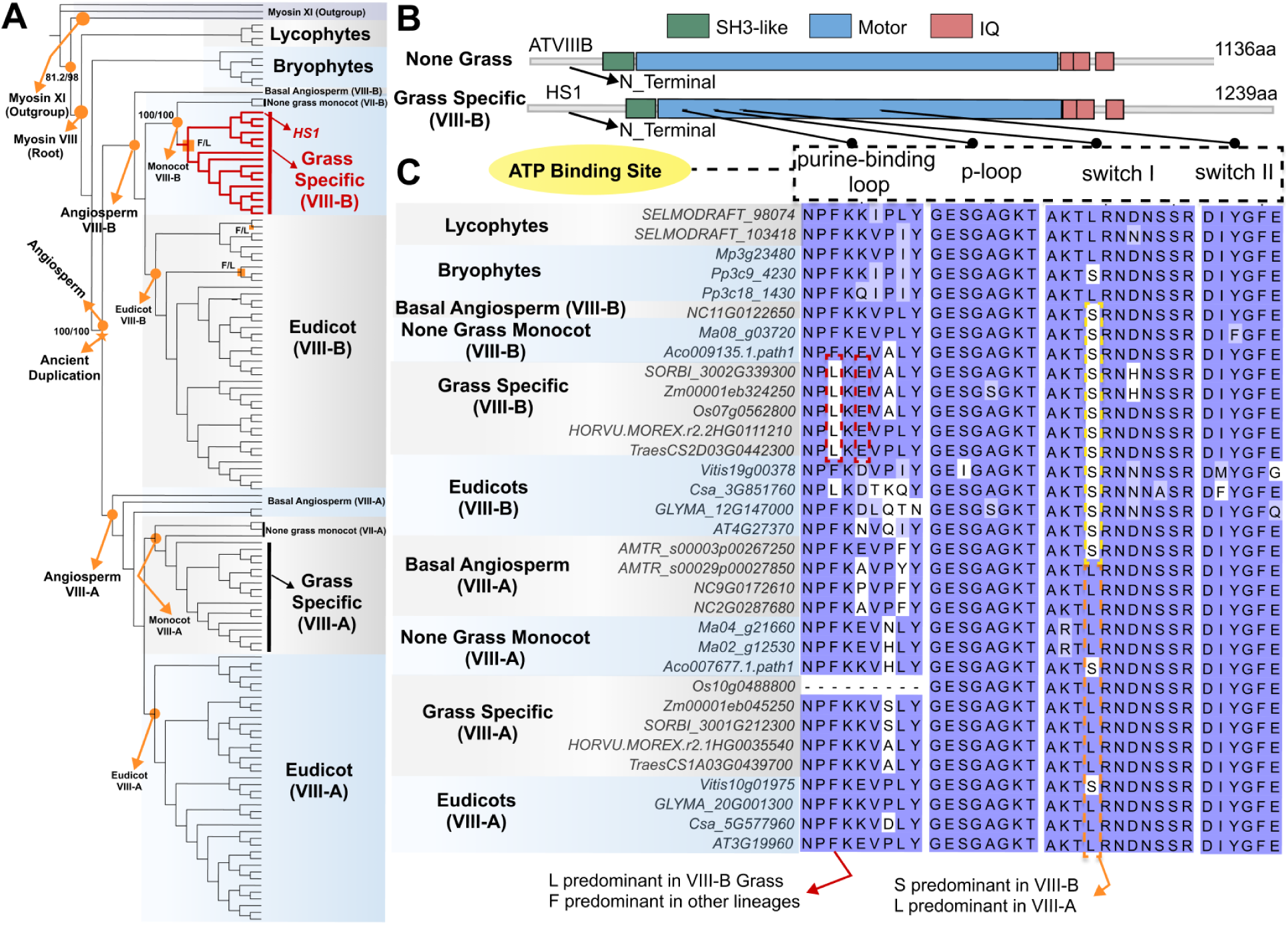
Phylogeny analysis of myosin proteins in plants. **A)** The phylogenetic tree of myosin family in plants. Branch support values are shown as SH-aLRT test/bootstrap, the orange circle indicates the lineage of the node. The orange square indicates the F/L substitutions in class myosin VIII. **B)** Schematic representation of the protein domain architecture of non-grass myosin VIII (*Arabidopsis thaliana* myosin VIII-B [ATVIIIB] as representative) and grass-specific myosin VIII-B (HS1 as representative). The N-terminal region is shown in gray; SH3-like (green), motor (blue), and IQ (red) domains are indicated. **C)** Multiple sequence alignment of the ATP-binding region of class VIII myosins from representative plant lineages. Lineage groupings are shown on the left. ATP-binding sites (purine-binding loop, P-loop, switch I, and switch II motifs) are indicated above the alignment. Conserved residues are shaded in blue. Dashed boxes highlight lineage-specific sequence features.

Accordingly, angiosperm myosin VIII proteins were grouped into four major clades (each represented by >10 species) for subsequent analyses: grass specific VIII-A, grass specific VIII-B, eudicot VIII-A, and eudicot VIII-B (Fig. 5A). Among these clades, the grass specific VIII-B clade exhibited the highest IUPred2 scores at both the N- and C-terminal regions (peak_N and peak_C = 0.81) (Supplementary Fig. S16A; Supplementary Data Set 2) and the longest median N-terminal intrinsically disordered region (IDR; 130.5 aa) (Supplementary Figs. S16, A and B; Supplementary Data Set 3), together indicating a strong propensity for intrinsic disorder (Mészáros et al., 2018). Notably, the grass specific VIII-B clade also displayed the longest median N-terminal length (181.5 aa) and uniquely retained both Motif 1 and Motif 2 (present in >90% of clade members), whereas other clades conserved only a single motif (Fig. 5B; Supplementary Figs. S17, A to C). Collectively, these features suggest that grass specific VIII-B myosins possess extended and disordered N-terminal regions that may facilitate dynamic protein–protein interactions or regulatory functions.

In addition to terminal disorder, grass specific VIII-B myosins exhibited lineage-specific substitutions within the motor domain (Figs. 5, B and C). Within the conserved purine-binding loop (NPxxxxxY), a leucine (L) at the third position was consistently present in grass specific VIII-B myosins, as well as in cucurbit species and Kalanchoe fedtschenkoi, whereas all other plant lineages retained phenylalanine (F) (Fig. 5; Supplementary Data Set 4). This L residue was consistently paired with a downstream glutamate (E), forming an L–E combination exclusive to grass specific VIII-B myosins (Fig. 5C; Supplementary Data Set 4). In addition, a leucine residue was uniquely conserved within the relay loop across grass specific VIII-B (Supplementary Fig. S18). These features were absent from non-grass myosin VIII proteins.

Structural modeling further revealed that *HS1* exhibited an estimated free energy of ATP binding of 2.267 kcal/mol, compared with 18.338 kcal/mol for *Arabidopsis* myosin VIII-A ATM1 (*AT3G19960*) and 6.677 kcal/mol for soybean myosin VIII-B (*GLYMA_12G147000*), indicating more favorable ATP binding in *HS1*, with approximately 8.1-fold and 3.0-fold lower ΔG values relative to ATM1 and soybean VIII-B, respectively (Supplementary Fig. S19). The F to L substitution occurs within the conserved purine-binding loop, a core component of the ATP-binding pocket (Risal et al., 2004), whereas other ATP-binding residues remained conserved across myosin VIII groups (Fig. 5C). These lineage-specific substitutions likely influence nucleotide binding, suggesting selective modifications of motor mechanics or grass-specific regulation.

Together, these lineage-specific features, including grass-specific N-terminal intrinsic disorder and motor-domain substitutions, indicate a distinct evolutionary trajectory for *HS1* that coincides with the divergence of *Poaceae* from other monocots.

### SCW-associated genes upregulated in *hs1* developing leaf

Analysis of publicly available transcriptome datasets (Goodstein et al., 2012; Papatheodorou et al., 2018) revealed that *HS1* was highly expressed (Fragments Per Kilobase of transcript per Million mapped reads, FPKM > 30) in sorghum leaves, inflorescences, and roots across multiple vegetative and reproductive stages, suggesting a broad role for *HS1* in sorghum development (Supplementary Fig. S20). To investigate transcriptome changes specifically in the developing leaf of the hs1 mutant, we performed RNA-seq on the hs1 line, which carries a nonsense mutation introducing a stop codon at amino acid position 573. Leaf base samples of developing and mature leaves with three biological replicates were collected at the sorghum GPD stage in the morning under optimal temperature and irrigation (Fig. 6A).

**Figure 6.**
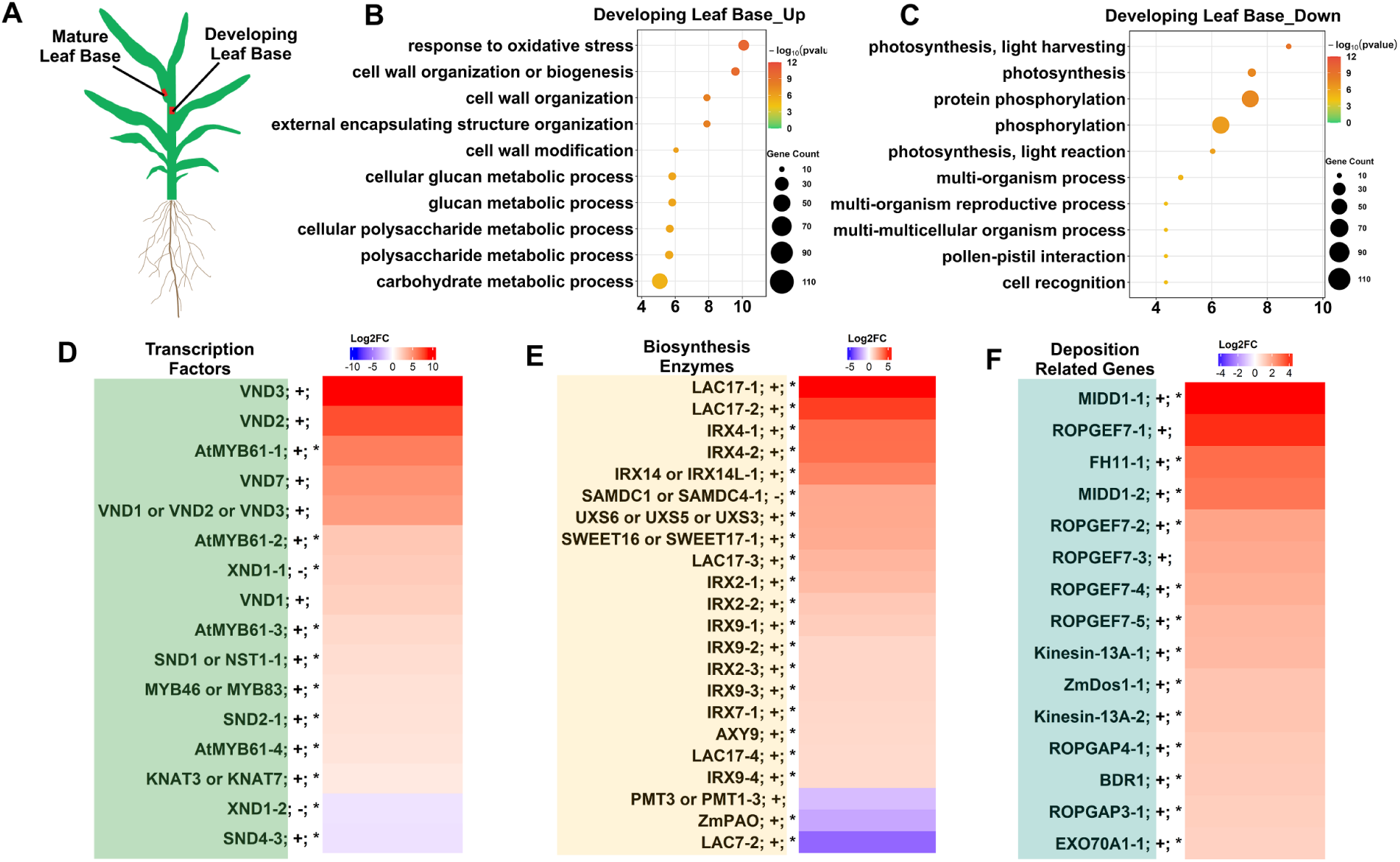
Transcriptome analysis of the midrib at the base of developing leaves in WT and *hs1*. **A)** The sketch map of tissues used in transcriptome analysis. **B, C)** GO enrichment analysis of upregulated (B) and downregulated (C) genes in developing leaf base midrib of *hs1* compared with WT. The top ten GO terms (P < 0.01), ranked by p-value. **D to F)** Heatmaps showing log_2_ fold changes in sorghum ortholog expression in *hs1* relative to WT for known secondary cell wall (SCW) related genes from *Arabidopsis thaliana*, *Zea mays*, and *Oryza sativa*, grouped as transcription factors (D), biosynthesis enzymes (E), and deposition-related genes (F); genes with |log2 fold change| > 1 are shown. “+” and “–” indicate positive and negative regulators of SCW formation, respectively. \**q*-value <0.05. The complete dataset is provided in Supplementary Fig. S20 and Supplementary Dataset 5.

Compared with WT, a total 2,383 upregulated and 1,242 downregulated differentially expressed genes (DEGs) (|log₂FC| > 1) were identified in developing leaves of *hs1*, whereas only 729 upregulated and 522 downregulated DEGs (|log₂FC| > 1) were detected in mature leaves (Supplementary Figs. S21, A and B). Consistent with the developmental onset of the *hs1* phenotype, more DEGs were detected in developing leaves than in mature leaves. Top ten up-regulated GO terms in the developing leaf base were enriched in cell wall–associated processes, such as cell wall biogenesis, organization and modification (Fig. 6B). Five out of top ten down-regulated GO terms were enriched in photosynthesis processes (Fig. 6C). In contrast, up-regulated genes in the mature leaf were enriched in general metabolic processes, such as RNA biosynthesis and macromolecule biosynthesis (Supplementary Fig. S21C). In addition, down-regulated DEGs were associated with transmembrane transport in mature leaves (Supplementary Fig. S21D).

To investigate whether *HS1* interfaces with known regulatory networks controlling protoxylem SCW formation, we analyzed transcriptional profiles of 128 sorghum homologs of characterized xylem SCW pathway genes from *Arabidopsis thaliana*, *Zea mays*, and *Oryza sativa*. This set included 38 transcription factors (TF), 53 biosynthetic enzymes (EZ), and 37 deposition-related (DP) genes (Supplementary Data Set 5). Strikingly, 99 (77.34%) genes were upregulated in *hs1*, with 64 exhibiting statistically significant induction [false discovery rate (FDR) P < 0.05] (Figs. 6, D to F; Supplementary Figs. S22, A to C; Supplementary Dataset 5).

The coordinated upregulation of SCW-associated genes in *hs1* indicates a transcriptional response to defective vascular structure. Despite robust activation of upstream regulators and biosynthetic enzymes, protoxylem vessels in *hs1* failed to maintain structural integrity. We therefore hypothesize that *HS1* is not required for the transcriptional activation of SCW genes but is essential for the effective execution of wall assembly. These results position *HS1* downstream of the core transcriptional network, where it likely facilitates cytoskeletal or mechanical processes that translate SCW gene expression into functional protoxylem reinforcement during leaf development.

### *HS1* regulates secondary wall integrity through lignin-associated pathways

Myosins function as molecular motors that transport diverse cellular cargos, including components associated with cell wall biosynthesis (Zhang et al., 2019). To determine which cell wall–related processes are associated with *HS1*, we reanalyzed a published sorghum single-cell RNA-seq (scRNAdeq) dataset of leaf tissues (Swift et al., 2024). In this dataset, during first leaf development (0-48 hours after germination, HAG), *HS1* showed the highest tissue expression specificity in the xylem cells compared with other cell types (Fig. 7A). Within the xylem cells, the highest expression was detected at 12HAG, when the first leaf is rapidly developing, compared to other timepoints. However, its expression decreased when the first leaf had fully matured (48HAG) (Fig. 7B, Supplementary Data Set 6). In another single-cell dataset, *HS1* expression remained low in mature leaf xylem (D’Agostino et al., 2025). This temporal and cell-type–specific expression pattern places *HS1* at the onset of protoxylem differentiation in developing leaves. Consistent with this pattern, the three most highly expressed xylem SCW biosynthetic genes at 12 HAG were all associated with lignification, including *irregular xylem 4* (IRX4/CCR1; Sobic.002G146000), *laccase 7* (LAC7; Sobic.003G357500), and *irregular xylem 9* (IRX9; Sobic.009G026101). Expression of these genes declined as leaves matured (48 HAG) (Fig. 7B; Supplementary Data Set 7). Notably, of these three genes, *irregular xylem 4* (IRX4/CCR1; Sobic.002G146000) and *laccase 7* (LAC7; Sobic.003G357500) belong to the lignin-specific biosynthetic pathway (Vanholme et al., 2010). In *Arabidopsis*, *CCR1* encodes a cytosolic cinnamoyl-CoA reductase (CCR) functioning in the cytosol, whereas *LAC7* encodes a laccase that catalyzes monolignol polymerization in SCW; both are required for the lignification of xylem SCW (Laskar et al., 2006; Zhao et al., 2015). In our bulk RNA-seq data, *CCR1 (IRX4)* was significantly upregulated (log₂FC = 3.75), whereas *LAC7* showed a strong down regulation in *hs1* (log₂FC = −3.44) (Supplementary Data Set 4). These suggested that *HS1* was likely involved in lignin synthesis within the xylem of developing leaves.

**Figure 7.**
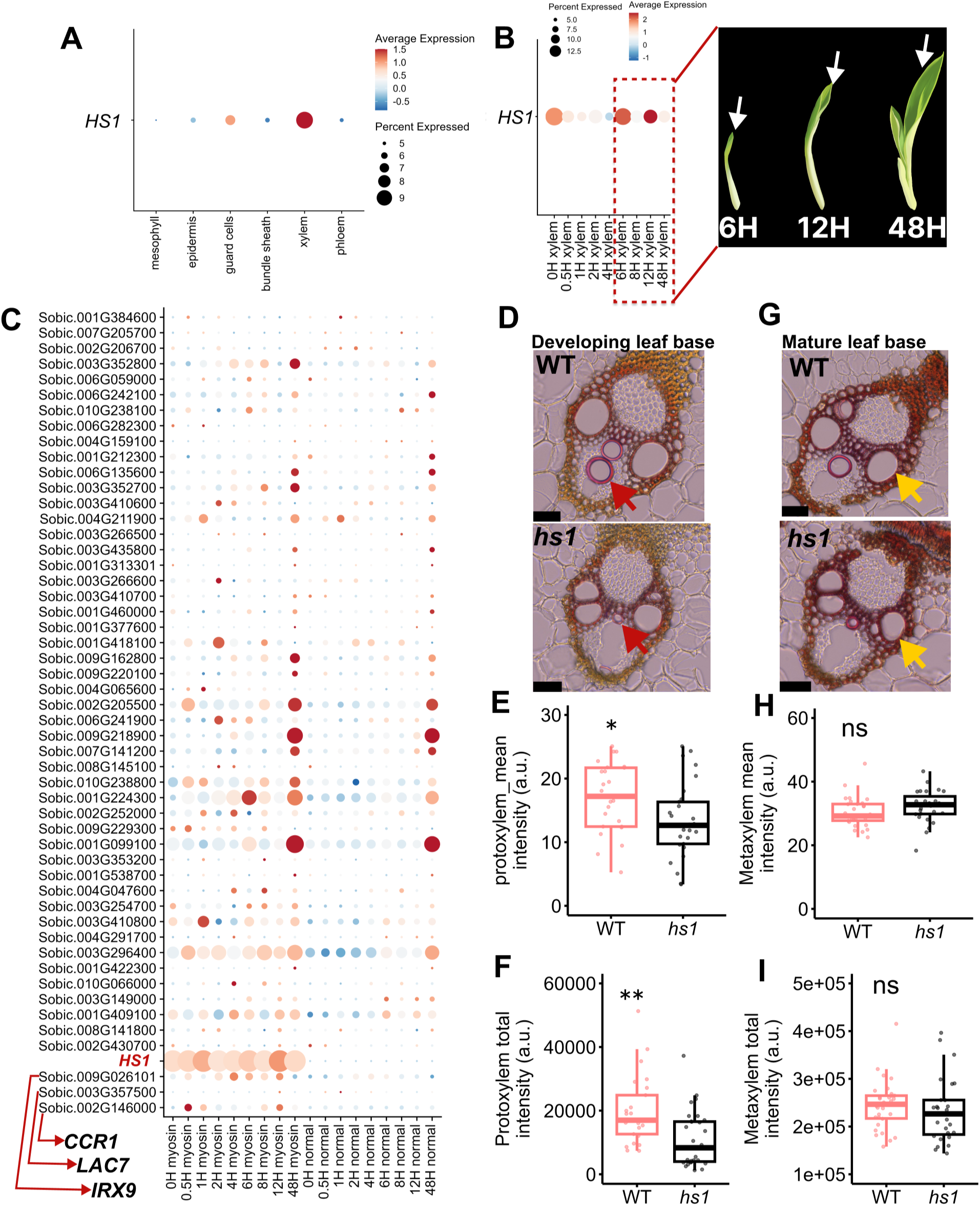
Coordinated single-cell expression of *HS1* and secondary cell wall (SCW) biosynthetic genes during leaf development and corresponding alterations in xylem lignification in *hs1* leaf midribs relative to WT. **A)** Dot plot of single-cell *HS1* expression across sorghum leaf cell types during first leaf development (0-48 hours after germination, HAG). **B**) Dot plot showing single-cell *HS1* expression in xylem cells across sorghum leaf developmental time points. The schematic at right depicts first-leaf morphology at 6 H (growth initiation), 12 H (rapid growth), and 48 H (maturation) (modified from Swift et al., 2024), with arrows indicating the first leaf. H, hours after germination. **C)** Dot plot of coordinated single-cell expression patterns of *HS1* and SCW biosynthetic genes in xylem cells across sorghum leaf developmental time points. Genes are ordered on the y-axis by average expression at 12 HAG, with the highest expression positioned at the bottom. X-axis labels indicate time points from 0 to 48 HAG. “Myosin” denotes xylem cells exhibiting high *HS1* expression, whereas “normal” denotes xylem cells exhibiting low *HS1* expression at the corresponding time points. Genes with high expression and patterns similar to *HS1* are indicated by red arrows. **D to I)** Comparisons of lignin intensity in developing and mature leaf base midribs of WT and *hs1*. Developing leaf bases: cross-sections (D), protoxylem (PX) lignin mean intensity (E), and total intensity (F). Mature leaf bases: cross-sections (G), metaxylem (MX) lignin mean intensity (H), and total intensity (I). Red and yellow arrows indicate protoxylem and metaxylem, respectively. Boxes indicate the first and third quartiles, and horizontal lines indicate the median. **P < 0.01; *P < 0.05; ns, not significant (P > 0.05); two-tailed Student’s t-test. Bars = 50 *µ*m. Data were collected from three biological replicates for each genotype. Quantification was performed on individual PX or MX vessels pooled across biological replicates (n ≥ 25 for D and E; n ≥ 27 for G and H).

To validate, we compared the lignin intensity in the xylem of developing leaves and mature leaves in WT and *hs1*. Lignin staining revealed visibly weaker signals in *hs1* protoxylem of developing leaves (Fig. 7C). Quantitative analysis revealed a significant reduction in *hs1* compared with WT, with a 21% decrease in mean lignin intensity and a 45% decrease in total lignin signal (P < 0.05 and P < 0.01, respectively; Figs. 7, D and E). In contrast, no significant difference (P > 0.05) was detected in mature leaves metaxylem SCW (Figs.7, F to H). Together, these results demonstrate that *HS1* is specifically required for proper lignin deposition during protoxylem SCW formation in developing leaves. The developmental specificity of the defect, combined with transcriptional mis-regulation of lignin polymerization genes, provides a mechanistic basis for protoxylem collapse and impaired water transport in *hs1*, positioning *HS1* as a critical regulator linking cytoskeletal function to lignin-mediated reinforcement of vascular cell walls.

### Low genetic diversity of *HS1* in the sorghum population

Given the essential role of HS1 in SCW development and protoxylem integrity, we next examined nucleotide diversity (π) at the *HS1* locus and its 20kb flanking regions in sorghum population. Nucleotide diversity were calculated using 400 Sorghum Association Panel (SAP) accessions (Bred, Conv, and Other) together with 46 wild sorghum (Wild) from the SorGSD database (Supplementary Data Set 8) (Morris et al., 2013; Liu et al., 2021; Boatwright et al., 2022). Notably, π was high across most of the flanking regions in all accessions but declined sharply in the *HS1* locus in both cultivated and wild sorghum (Fig. 8A). Across SAP subgroups, diversity at *HS1* was reduced by approximately seven-fold compared to the immediately adjacent 20-kb regions, consistent with strong local purifying selection or selective constraint acting on this gene. This pattern indicates that HS1 has experienced intense selective pressure in sorghum population, likely preserving its function in maintaining vascular robustness under the heat- and drought-prone conditions typical of sorghum-growing environments.

**Figure 8.**
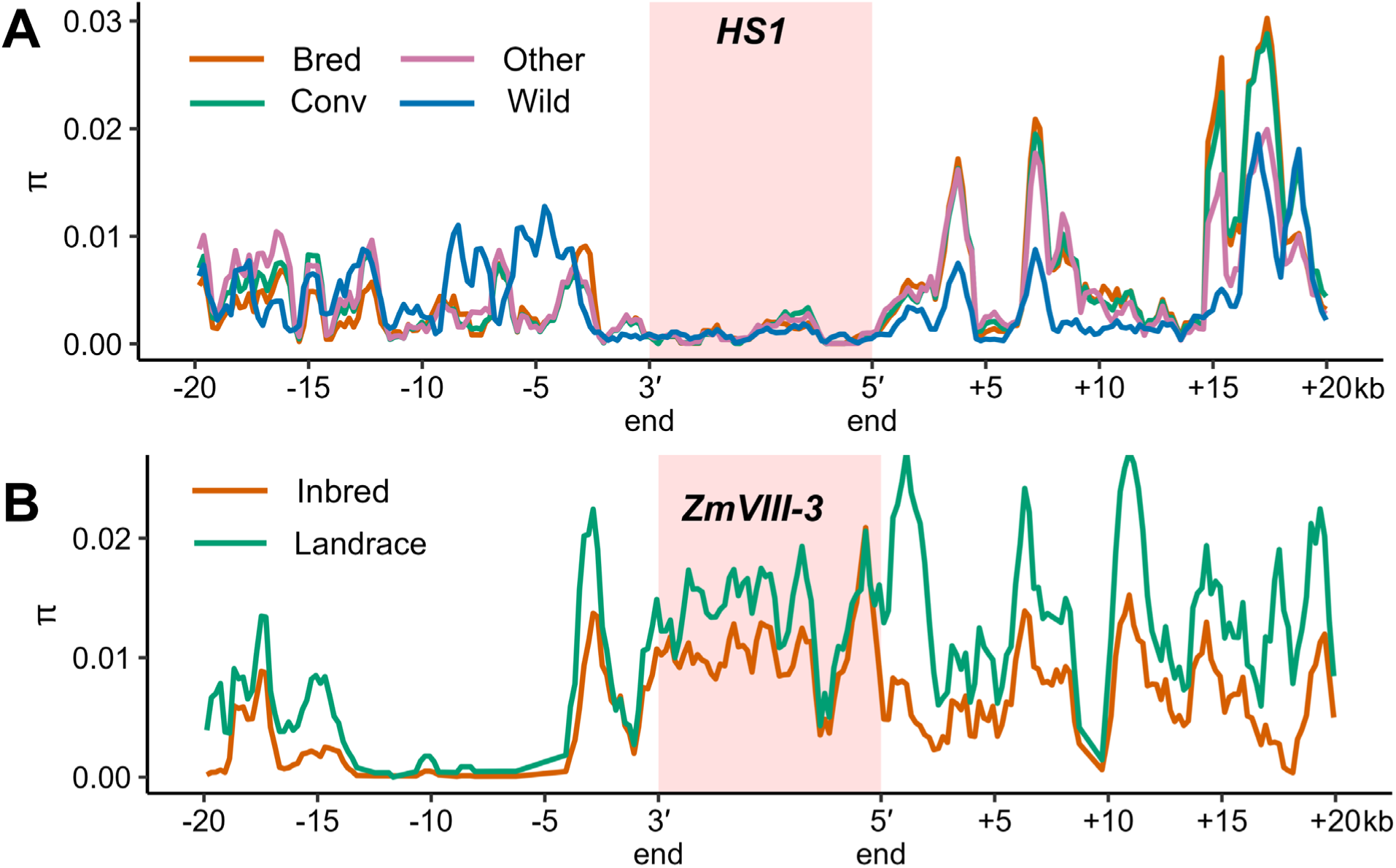
Evolutionary conservation profiles at the *HS1* and ZmVIII-3 loci. A) Nucleotide diversity (π) across the *HS1* locus and its 20-kb flanking regions in breeding lines (Bred), photoperiod-converted lines (Conv), wild accessions (Wild), and accessions without subgroup assignment (Other). The shaded region denotes the HS1 gene body. B) Nucleotide diversity (π) across *ZmVIII-3* and its 20-kb flanking regions. The shaded region denotes the *ZmVIII-3* gene body.

In striking contrast, the maize homolog at the *ZmVIII-3* (*Zm00001d021785*) locus exhibited the opposite pattern, with consistently high nucleotide diversity maintained across diverse maize populations (Fig. 8B). This suggests differential evolutionary trajectories for this conserved gene between the two species, possibly reflecting differences in selective pressures related to vascular development or environmental adaptation.

## DISCUSSION

### Potential Impact of *HS1* Discovery

Efficient vascular development is central to plant growth and environmental resilience, yet the molecular mechanisms that coordinate motor protein with SCW assembly in xylem development remain unclear. Here, we identify *HS1*, a grass-specific myosin VIII protein, as a previously unrecognized regulator of protoxylem SCW integrity that is essential for maintaining hydraulic function and heat tolerance during leaf development. Through forward genetics, identification of *HS1* highlights a critical divergence between Arabidopsis-based xylem paradigms and the vascular system of grasses. Unlike *Arabidopsis* and other dicots, grasses produce long, rapidly elongating leaves with high transpirational demand, thereby requiring protoxylem that remains mechanically stable and hydraulically functional during active growth (Scoffoni et al., 2017). Studying sorghum mutants allowed us to examine protoxylem differentiation within rapidly elongating leaves under sustained developmental and hydraulic context, largely absent from Arabidopsis systems (Roeder et al., 2025; Uauy et al., 2025). While extensive transcriptional regulators of SCW formation and xylem differentiation have been defined in Arabidopsis, regulators directly controlling leaf protoxylem SCW execution and water transport have remained elusive. *HS1* fills this knowledge gap by establishing a mechanistic framework linking a cytoskeletal motor protein to lignin-associated SCW reinforcement and stress resilience in sorghum. Notably, *HS1* is continuously required across successive leaves and vegetative stages to maintain protoxylem integrity and water transport (Supplementary Figs. 7, A and B). The strong conservation and remarkably low π in the *HS1* loci across sorghum populations, including wild accessions (Fig. 8A), further supports its designation as a core regulator in leaf protoxylem development.

Our work demonstrates, to our knowledge, that a motor protein can directly influence whole-plant physiological traits. Previous work on plant myosins and kinesins largely focused on cytoskeletal organization, organelle transport, or cell division (Reddy and Day, 2001b; Hussey et al., 2002; Smith, 2003; Sparkes, 2011; Nebenführ and Dixit, 2018). In contrast, our study suggests that *HS1* links intracellular motor activity to biomechanical wall reinforcement and vascular transport capacity. Safranin staining demonstrated that water transport along developing hs1 leaves was markedly slowed (Figs. 2, A to D), indicating restricted longitudinal hydraulic delivery due to protoxylem structural failure. In addition to the collapsed protoxylem morphologies observed ∼5 mm above the leaf sheath (Supplementary Fig. S6), serial transverse sections along the developing leaf base midrib (0–10 mm above the leaf sheath) further revealed progressive protoxylem deformation and collapse in *hs1* (Supplementary Fig S23). This likely contributed to the delayed internal water transportation likely underlies the scorching phenotype, as insufficient supply to expanding tissues renders them vulnerable under elevated evaporative demand (Fig. 9; Supplementary Figs. S9, A to D). Notably, recent work showed that α-tubulin–mediated alterations in microtubule organization similarly disrupt protoxylem wall patterning and reduce hydraulic conductivity, as evidenced by safranin transport assays (Huang et al., 2024). In contrast to the structural role of tubulin in microtubules, *HS1* operates as a force-generating motor on actin filaments that preserve protoxylem integrity. Together, these findings suggest that cytoskeletal regulation of protoxylem function operates at multiple hierarchical levels, from microtubule-guided pattern specification to motor-dependent wall execution. Within this regulatory framework, *HS1* exhibits distinctive evolutionary features.

**Figure 9.**
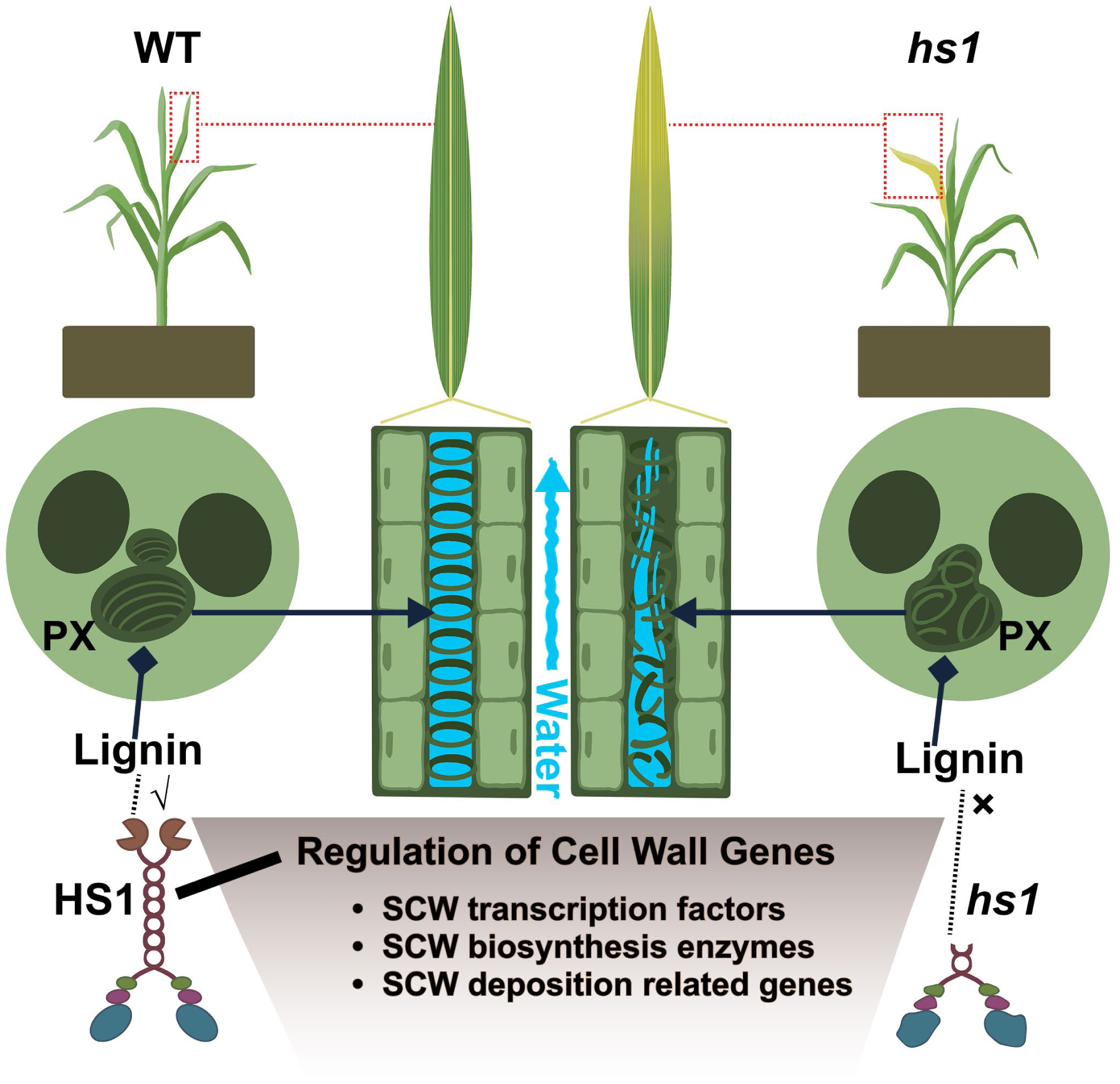
Proposed model for *HS1* function in maintaining protoxylem secondary cell wall (SCW) integrity. *HS1* regulates SCW gene expression and lignin deposition to maintain protoxylem structure and water transport. Loss of *HS1* leads to reduced lignin, protoxylem collapse and impaired water transport.

Lineage-specific features further indicate that *HS1* belongs to a grass-specific myosin VIII-B lineage and retains lineage-specific structural features, due to its extended intrinsically disordered N-terminal region and conserved VIII-B motifs (Supplementary Fig S16; Supplementary Fig S17). These features likely enable protein–protein interactions essential for protoxylem SCW deposition and xylem function. Structural modeling predicts reduced ATP-binding free energy for *HS1*, potentially indicating enhanced motor activity. Together, these findings raise the possibility that lineage-specific structural features and altered motor properties contributed to functional specialization during vascular evolution.

Consistent with this evolutionary specialization, our integrated analyses support a hierarchical A–B–C organization of SCW assembly, in which upstream biosynthetic supply modules (A) and downstream wall-localized execution factors (C) are coordinated through a central execution node (B). *HS1* occupies this intermediate B layer. In *hs1*, induction of modules A fails to compensate for reduced C-layer execution factors (Supplementary Data Set 9), resulting in defective wall assembly. The presence of multiple MYB-binding sites in the *HS1* promoter, along with altered expression of sorghum MYB homologs in *hs1*, supports the positioning of *HS1* downstream of the MYB transcriptional tier within the proposed A–B–C framework (Supplementary Data Set 10; Supplementary Data Set 11). Developmental and single-cell expression analyses further refined this framework by demonstrating that *HS1* expression is enriched in differentiating xylem cells and is specifically associated with protoxylem development. Together, our findings identify *HS1* as a cytoskeleton-based execution hub that links SCW gene activation to the physical construction of lignified vessel walls in protoxylem. This work expands the current SCW regulatory paradigm beyond transcriptional control and highlights myosin proteins as previously unrecognized determinants of vascular integrity and hydraulic function, opening new avenues for investigating cytoskeleton-dependent mechanisms in plant vascular development.

Finally, the developmentally sustained impact of *HS1* establishes clear agricultural relevance. Loss of *HS1* disrupts protoxylem integrity, triggering a cascade of hydraulic and physiological consequences, from leaf scorching to impaired transpiration and diminished photosynthetic capacity. Importantly, these effects persist across developmental stages and multiple leaves. This underscores the central importance of xylem integrity for plant performance and positions *HS1* as a promising target for future crop improvement. Leveraging *HS1* function or natural allelic variation may provide novel routes to improve water transport and drought tolerance, supporting sustainable crop production in thermally and hydrologically challenging environments. The discovery of *HS1* therefore bridges fundamental plant biology with translational opportunities in grass crops.

### Outstanding Questions and Future Directions

While *HS1* established a functional framework for motor protein activity in protoxylem, several mechanistic questions remain. Foremost is the identity of *HS1*’s interacting cargo and interaction partner and how motor-driven transport contributes to SCW assembly. In Arabidopsis, classical irregular xylem (irx) mutants have defined a core set of secondary cell wall biosynthetic enzymes (Turner and Somerville, 1997; Taylor et al., 2000; Jones et al., 2001; Taylor et al., 2003; Brown et al., 2005; Brown et al., 2007; Peña et al., 2007; Persson et al., 2007; Brown et al., 2009) embedded within MYB-directed transcriptional networks (Zhong et al., 2007; Zhou et al., 2009; Zhong and Ye, 2015; Kumar et al., 2016), outlining a molecular landscape that may serve as potential cargo for motor-driven transport. Identifying proteins that physically interact with *HS1* and defining its direct cargo during protoxylem development are the next crucial steps. This will likely require defining *HS1*-associated complexes (e.g., affinity purification–mass spectrometry), validating candidate interactions, and directly testing *HS1*-dependent trafficking by live-cell imaging, followed by genetic perturbation and epistasis analyses of top candidate cargos.

Additional questions concern the mechanistic basis of lineage-specific adaptations. *HS1* belongs to the myosin VIII family, which remains comparatively understudied relative to myosin XI (Ryan and Nebenführ, 2018). Grass specific VIII-B proteins are distinguished by extended N-terminal intrinsically disordered regions and conserved lineage-specific motifs. How these structural features modulate motor activity and contribute to protoxylem SCW formation, hydraulic function, and stress adaptation warrants further investigation.

Finally, translating *HS1* knowledge to crops presents opportunities and challenges. Whether natural or engineered HS1 alleles can enhance xylem efficiency and water-use traits across grass remains an open question. For example, *ZmVIII-3*, the maize homolog of *HS1*, exhibits substantially higher sequence diversity across populations and lacks the extended tailed and coiled-coil helix compared to *HS1* (Fig. 8A; Supplementary Fig. S24). This divergence suggests that maize may harbor unexplored natural variants with *HS1*-like architectures, while also pointing to protein structural remodeling as a plausible translational direction. More broadly, these observations raise the possibility that lineage-specific structural innovation may represent a recurrent strategy for optimizing vascular function in grasses. This perspective leads to a broader question: are other plant motor proteins similarly co-opted for vascular or stress-related functions? Answering these questions will clarify both the evolutionary trajectory of motor protein function and its potential for crop improvement.

## MATERIALS AND METHODS

### Identification of the causal mutation of *hs1*

The *hs1* mutant was identified through field screening of pedigreed M₃ mutant populations generated from EMS-treated BTx623 (Jiao et al., 2016) at the USDA research fields in Lubbock, Texas (33°41′36.4596″ N, 101°54′18.612″ W; elevation 992 m). Field trials were conducted under fully irrigated conditions using a subsurface drip irrigation (SDI) system that delivered approximately 1 inch of water per week.

To map the causal mutation, the *hs1* mutant was crossed with BTx623 to produce F₁ progeny. F₁ plants were self-pollinated to generate F₂ populations, which were evaluated for leaf scorching phenotypes under field conditions during the 2018 and 2021 growing seasons.

Identification of the causal mutation was performed using a previously established BSA pipeline (Jiao et al., 2018a; Jiao et al., 2018b; Wang et al., 2021). Briefly, genomic DNA was extracted from leaf tissue of 20 F₂ individuals exhibiting the mutant phenotype, using a CTAB-based protocol as described previously (Xin and Chen, 2012). DNA samples were quantified, pooled in equal proportions, and used to construct an Illumina paired-end (150 bp) library, yielding an average genome coverage of ∼12–15×. Sequencing reads were processed through our established workflow (Jiao et al., 2018b) and aligned to the *Sorghum bicolor* BTx623 V3 reference genome (McCormick et al., 2018) with Bowtie2 (Langmead and Salzberg, 2012). Variant calling was done using BCFtools (Narasimhan et al., 2016). To specifically capture EMS-induced mutations, SNPs were filtered to retain only GC→AT transitions, exclude background polymorphisms identified in resequencing data from our BTx623 germplasm, and satisfy read depth (5–100) and homozygosity criteria. Functional impacts of filtered variants were annotated using SnpEff (Cingolani et al., 2012). Candidate causal mutations were defined as variants predicted to cause amino acid substitutions, premature stop codons, or splice-site alterations. To assess allelic recurrence, candidate variants were queried against an indexed sorghum mutant database (Jiao et al., 2024).

### Physiological characterization of *hs1* under a controlled environment

For all controlled-environment experiments, WT and *hs1* sorghum plants were cultivated in a phytotron greenhouse (Lubbock, TX, USA) under standard irrigation. Plants were maintained at 28/25 °C (day/night) with a 16-h light/8-h dark photoperiod. Due to diurnal fluctuations, the actual greenhouse temperature ranged from approximately 26 to 30 °C, within the optimal range for sorghum growth. All experiments below are conducted at the GPD stage (∼30 days after planting).

#### Root

Whole root systems were gently removed from the pots. Adhering soil was washed away under running tap water with great care to avoid mechanical damage and to preserve fine lateral roots. Cleaned roots were spread on black cardboard and photographed. The resulting images were analyzed by ImageJ (Schneider et al., 2012) for root morphology and root surface area. For sap exudation measurements, plants at the same developmental stage were cut 2 cm above the soil surface. To measure xylem sap exudation under minimally disturbed conditions, plants were maintained under near-saturated humidity to minimize transpiration and evaporation. Stems were cut above the stem base, and the cut end was immediately covered with an inverted 50-mL centrifuge tube to maintain high humidity. Xylem sap was allowed to exude naturally without physical contact or applied pressure. Exuded sap was gently collected using pre-weighed absorbent paper, and the paper was reweighed immediately. Sap exudation was calculated as the increase in paper weight in two hours.

#### Stomata

To obtain leaf epidermal impressions, a thin layer of clear nail polish was applied to the abaxial surface of developing leaves, allowed to dry, and lifted with transparent tape onto a glass slide. Stomatal images were captured using an EVOS M5000 microscope system. Stomatal density and stomatal pore length were quantified in ImageJ (Schneider et al., 2012). For each plant, stomatal density was calculated from at least three images. Stomatal size (represented as guard cell length) (Jordan et al., 2015) was measured from 20 stomata per plant. Three biological replicates were analyzed for each genotype.

#### Water transport

For above-ground tissue assays, WT and *hs1* plants were cut at the base of the stem, and the entire above-ground portion was placed into 0.1% safranin solution on a sunny day. The total weight of the container, solution, and excised tissue was recorded at the start and again after 2 h to quantify dye uptake. For leaf assays, developing leaves were excised at the base and immersed in 0.1% safranin solution under the same conditions. The total weight was measured at the beginning and after 10 min to calculate dye absorption and movement. Three biological replicates were analyzed for each genotype.

Water-loss assay: developing leaves were detached from WT and *hs1* plants, with three biological replicates per genotype. Samples were placed at room temperature and weighed at 15, 30, and 45 min, and at 1 and 2 h. Water loss was calculated and expressed as the percentage of the initial fresh weight. Three biological replicates were analyzed for each genotype.

Leaf physiological indices: developing leaves from at least three biological replicates were used for each genotype. The H_2_O_2_, leaf relative water content (LRWC), total chlorophyll (Chl) and electrolyte leakage (EL) assays were performed as previously described with minor modification (Mostofa et al., 2015; Mostofa et al., 2020). Briefly, to detect H_2_O_2_ accumulation in situ, samples were freshly cut and transffered to 0.1% 3,3-diaminobenzidine (DAB) for 72h and then were bleached in acetic acid-glycerol-ethanol (1/1/3) (v/v/v) solution for 24h and then photographed. LRWC was measured to assess leaf water status. Fresh leaves were cut and immediately weighed to obtain fresh weight (FW). Samples were fully hydrated by submerging them in deionized water for 24 h in 50-mL tubes placed on a horizontal shaker, after which turgid weight (TW) was recorded. Leaves were then dried at 70 °C for 72 h to obtain the dry weight (DW). LRWC was calculated as: LRWC (%) = [(FW − DW) / (TW − DW)] × 100. To determine Chl content, 0.1 g leaves were homogenized in 2 mL chilled 80% acetone and centrifuged at 11,500 × g for 12 min at 4 °C. Absorbance of the clarified supernatant was measured at 663, 645, and 470 nm using 80% acetone as the blank. Total Chl was calculated using previously established formula (Arnon, 1949) :

Total Chl content (mg g⁻¹ FW) = (20.2 × A645 + 8.02 × A663) × V / (1000 × W)

where V denotes the volume of 80% (v/v) acetone used for extraction (mL), and W represents the fresh weight of the tissue sample (g). EL was assessed by incubating ∼0.3 g of 5-cm leaf segments with the midrib removed in 45 mL deionized water at room temperature on a horizontal shaker for 24 h. The conductivity of the bathing solution (C₁) was then measured, and the tubes were boiled for 30 min to release total electrolytes and obtain the final conductivity (C₂). EL was calculated as (C₁ / C₂) × 100.

#### Stomatal conductance

WT and *hs1* plants stomatal conductance and E_apparent of the developing leaves (9^th^ leaf) were measured with an LI-600 porometer (LI-COR, USA). Measurements were conducted in the morning and again near noon (before the onset of leaf scorching symptoms). Five biological replicates were analyzed per genotype, with three measurements taken per replicate and averaged.

### Scanning electron microscopy

Fresh WT and *hs1* samples were collected from the bases of developing leaves at the five-leaf, GPD, and flag-leaf stages. Samples were collected at night to reduce transpiration-driven xylem tension. Approximately 2-mm transverse sections were excised and immediately immersed in liquid nitrogen, followed by transfer to a freeze dryer (Model 117, Labconco Corporation). After overnight lyophilization, dried sections were mounted onto aluminum stubs for imaging. Scanning electron microscopy was performed using a Hitachi S-3400 microscope operated at an accelerating voltage of 20 keV. Lumen area, long axis, short axis, and SCW thickness of protoxylem vessels were measured manually using ImageJ (Schneider et al., 2012). Three biological replicates were analyzed for each genotype.

### Transcriptome profiling of leaf base midrib

At the GPD stage (∼30 days after planting), WT and *hs1* plants were grown in the greenhouse under the conditions described above. Total RNA was extracted from the midrib of the base of mature and developing leaves (the one just above the last fully expanded mature leaf), using the RNeasy® Plant Mini Kit (74904; QIAGEN) according to the manufacturer’s instructions. RNA sequencing libraries were prepared using standard Illumina protocols and sequenced by a commercial service provider. For each sample, three independent biological replicates were sequenced.

Clean RNA-seq reads were mapped to the Sorghum bicolor BTx623 reference genome (version 3) using STAR (Dobin et al., 2013). Gene expression levels for each biological replicate were estimated as FPKM (fragments per kilobase of transcript per million mapped reads) using Cufflinks (Trapnell et al., 2012). Reproducibility among biological replicates was evaluated by calculating Pearson correlation coefficients in R. To ensure data reliability, two replicates were excluded from downstream analyses because their coloration was below 0.95 when compared with other corresponding replicates. Differentially expressed genes (DEGs) were identified using the CuffDiff module of the Cufflinks package (Trapnell et al., 2012). Functional enrichment analysis of DEGs was performed using agriGO Singular Enrichment Analysis with a hypergeometric test, applying a significance threshold of P < 0.01 (Tian et al., 2017). Samples were collected with three biological replicates for each genotype.

### Identification of orthologs

We conducted BLAST searches to determine the homologous relationships between sorghum and those xylem development genes characterized in *Arabidopsis*, *Zea mays*, *Oryza sativa*. Query sequences were aligned against the sorghum BTx623 genome (V3.1). Our criteria for identifying similarity encompassed the first five hits exhibiting a percentage similarity exceeding 30%, an E-value of 1e-5, and a score cut-off of 100. Subsequently, reverse BLAST was performed using the top hits against the sorghum protein sequence. If the same sorghum protein, initially queried in the context of rice, Arabidopsis, and *Z. mays*, emerged among the top three hits within the sorghum amino acid sequence, it would indicate a predicted orthologous relationship between the respective genes.

### Phylogenetic analysis

Two phylogenetic analyses were conducted to classify *HS1* as myosin VIII or XI and to determine its phylogenetic position within the myosin VIII clade. For the classification analysis (Supplementary Fig. S13), myosin VIII and XI sequences from Arabidopsis, maize, rice, and sorghum were included. Arabidopsis and maize sequences were obtained from published datasets (Reddy and Day, 2001a; Wang et al., 2014). Rice and sorghum sequences were retrieved from RAP-DB (Kawahara et al., 2013; Sakai et al., 2013) and Phytozome (Goodstein et al., 2012), respectively. Sequences were aligned using ClustalW, and a neighbor-joining tree was constructed in MEGA 11.0.13 under the p-distance model with pairwise gap deletion and 1,000 bootstrap replicates (Tamura et al., 2021; Madeira et al., 2024); trees were visualized in FigTree (Rambaut, 2014).

To determine the phylogenetic position of *HS1* within myosin VIII (Fig. 5A), 134 myosin VIII homologs from 44 land species were retrieved from Ensembl Plants via BioMart (Kinsella et al., 2011; Yates et al., 2022) by filtering motor domain (MYSc_Myo8). Full-length sequences were aligned using the same method. Maximum-likelihood inference was performed with IQ-TREE v3.0.1 using the ModelFinder-selected substitution model Q.PLANT+I+R6 under BIC (Kalyaanamoorthy et al., 2017; Wong et al., 2025). Branch support was evaluated with SH-aLRT and ultrafast bootstrap tests (1,000 replicates each) (Hoang et al., 2018). Three Arabidopsis myosin XI proteins served as outgroups (Supplementary File 1). rees were visualized in iTOL (Letunic and Bork, 2024)

### Terminal feature analysis of myosin VIII proteins

Protein domain architecture was determined using InterProScan (Jones et al., 2014) with ProSiteProfiles (Sigrist et al., 2026) annotations. Based on ProSiteProfiles-defined domain boundaries, the N-terminal region was defined as the sequence preceding the first detected SH3 domain, while the C-terminal boundary was defined by the position of the last detected IQ motif and was used solely to delineate conserved domain architecture.

In N terminal, sequence motif patterns were analyzed using the MEME suite (Bailey et al., 2015), and motif patterns were plotted using the TBtools (Chen et al., 2020). Intrinsic disorder (IDR) of myosin VIII proteins was analyzed using IUPred2A (Mészáros et al., 2018) in long-disorder mode.For each protein, residue-wise IUPred2 scores were calculated across the full-length sequence. To generate group-level disorder profiles, residue positions were normalized by protein length and binned into 100 equally sized relative-position bins along the protein. Mean IUPred2 scores were calculated for each bin within each group, producing averaged disorder profiles that reflect lineage-specific trends in intrinsic disorder distribution. Comparative analyses were restricted to groups containing more than ten species to ensure sufficient sampling depth.

For visualization, normalized residue positions were mapped onto the length of a representative reference protein (*HS1*), allowing direct comparison of IDR profiles relative to conserved domain structure. N terminal IDR were quantified by counting residues with IUPred2 scores ≥ 0.5 within the N-terminal region.

### Motor domain feature analysis of myosin VIII proteins

Conserved functional features within the myosin motor domain were analyzed using a reference-guided, feature-centred alignment strategy. Motor domain boundaries were first identified based on Pfam annotations, and the extracted motor-domain sequences were aligned using a using hmmalign in HMMER v3.1b2 (Eddy, 2011).

Feature regions within the motor domain, including the purine-binding loop, P-loop, switch I, switch II, relay loop, and a downstream converter-region window, were defined as fixed amino acid intervals in a representative reference protein (*HS1*). Reference residue positions were mapped to alignment columns by sequentially indexing ungapped residues in the reference sequence within the multiple sequence alignment, followed by manual verification. For each feature, aligned residue blocks corresponding to the defined column ranges were directly extracted from all sequences, retaining alignment gaps to preserve positional correspondence across species. Feature-specific alignments were used for residue-level comparative analyses.

For visualization, feature-specific alignments (Supplementary Data Set 4) were examined in Jalview (Waterhouse et al., 2009), and a subset of sequences representing the diversity within each phylogenetic group was selected for display in the main figures. To summarize residue conservation patterns at the population level, sequence logos were generated for each feature using all aligned sequences.

ATP docking analysis was performed using ATPdock (Rao et al., 2022). First, ATP binding sites for previously generated PDB files (see Protein structure and stability analysis) were obtained using ATPbind (Hu et al., 2018). The PDB files along with the binding site information were analyzed using ATPdock with default parameters and the estimated free energy was obtained for each protein. ATP pocket visualization was performed with PyMOL (Schrodinger, 2015).

### Single cell RNAseq analysis

The published single cell data from Swift et al. (2024) was obtained from the NCBI gene expression omnibus (GSE248919) for further analysis. Single cell raw reads from D’Agostino et al. (2025) was obtained from the authors and processed as detailed in the manuscript. Analysis was performed in R using the Seurat package (version 4.4) (Hao et al., 2021). Cells with no assigned cell type were removed before further investigation. Xylem cells were subset from the whole dataset and grouped into two groups based on *HS1* gene expression. Cells with > 0 normalized expression of hs1 were labeled as “myosin” and cells with <= 0 were labeled as “normal”. Gene expression was visualized using the dotplot function in the Seurat package.

### Histological analysis

At the GPD stage (∼30 days after planting), fresh midrib tissues from the same basal region of mature and developing WT and *hs1* leaves were collected at night to reduce transpiration-driven xylem tension and embedded in Tissue Plus O.C.T. Compound (Fisher HealthCare, Cat# 4585, Houston, TX, USA). Samples were transversely sectioned at ∼30 µm thickness using a cryostat. Fresh sections were transferred onto charged slides and rinsed with water for 2 min to remove residual O.C.T. The slides were then incubated in 2% phloroglucinol prepared in 95% ethanol for 5 min. Concentrated HCl was applied to induce the Wiesner reaction, and stained tissues were imaged immediately using a light microscope (Invitrogen EVOS M5000 Imaging System) under identical acquisition settings. Lignin deposition was quantified in ImageJ (Schneider et al., 2012) by measuring the mean and total staining intensity of protoxylem SCW in developing leaf-base midribs and of metaxylem SCW in mature leaf-base midribs, with all measurements corrected by background subtraction. Samples were collected with three biological replicates for each genotype.

### Nucleotide diversity analysis

To investigate the evolutionary patterns of myosin across species, we quantified nucleotide diversity (π) at the *HS1* locus, and its maize ortholog ZmVIII-3 (*Zm00001d021785*), together with the 20-kb regions flanking each gene. Genome-wide sorghum variants for 400 Sorghum Association Panel (SAP) accessions were obtained from the European Variant Archive under accession PRJEB51985 (Morris et al., 2013; Boatwright et al., 2022). Variant data for wild sorghum were retrieved from the the Sorghum Genome SNP Database (SorGSD) for wild sorghum (Liu et al., 2021). Maize inbred and landrace variant datasets were obtained from the unified high-coverage VCF dataset released by MaizeGDB (Andorf et al., 2025), based on the B73 RefGen_v5 reference genome. π values were calculated by VCFtools (v0.1.17) within 800-bp windows with a 200-bp step across the region (Danecek et al., 2011).

## Supporting information

Supplementary Data Sets

## DATA AVAILABILITY

All sequencing data have been deposited in the NCBI SRA database, including transcriptome data (BioProject PRJNA1419276) and whole-genome genomic DNA sequencing data of the mutants (SRR18888791, SRR18888802, SRR18888804, and SRR18888806).

## ACKNOWLEDGMENTS

We thank Drs. Lam-Son Phan Tran and Zhixin Xie from Texas Tech University for their valuable comments and suggestions for this project.

## AUTHOR CONTRIBUTIONS

Y.J., G.P., and J.C. conceived the project. J.C., Y.J., Z.X., and T.R. identified the *hs1* mutant and mapped the causal gene. Z.L. and T.R. conducted physiological experiments. Z.L., T.R., and L.D. performed the data analysis. Z.L., T.R., Y.J., and G.P. drafted the manuscript. All authors reviewed and edited the manuscript and approved the final version.

