## Supplementary Figures for "A Grass-Specific Structural Feature of Myosin VIII Regulates Protoxylem Development and Hydraulic Conductance in Sorghum"

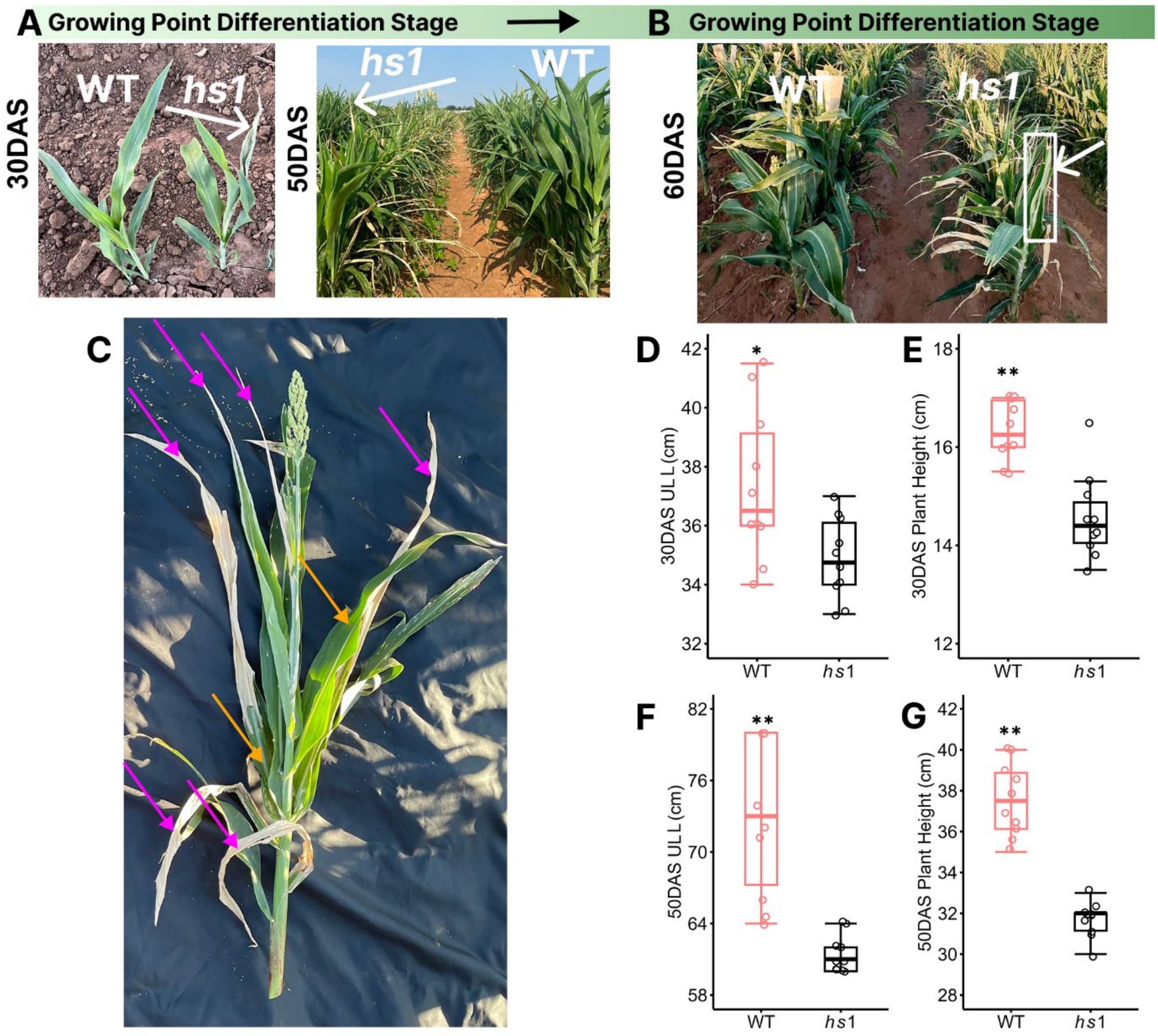


**Supplementary Figure S1.** Phenotype of wild type (WT, BTx623) and *hs1* in the field. **A, B)** Leaf scorching phenotype of WT and *hs1* in a sorghum field in Lubbock, TX during summer during 30, 50 and 60 days after sowing (DAS); white arrows point to scorching developing leaf at growing point differentiation (GPD) (A) and flag leaf (B) stage**. C)** heat wave–induced damage in *hs1* leaves. Magenta arrows highlight scorched upper leaf regions, while orange arrows indicate the green basal regions. **D** to **G)** Comparisons of upper leaf length (ULL) and plant height between WT and *hs1* at 30 and 50 DAS. (D) ULL at 30 DAS; (E) plant height at 30 DAS; (F) ULL at 50 DAS; (G) plant height at 50 DAS. For all statistical analyses, boxes indicate the first and third quartiles, horizontal lines denote median, n = 10. **P < 0.01, *P < 0.05 (two-tailed Student's t-test).

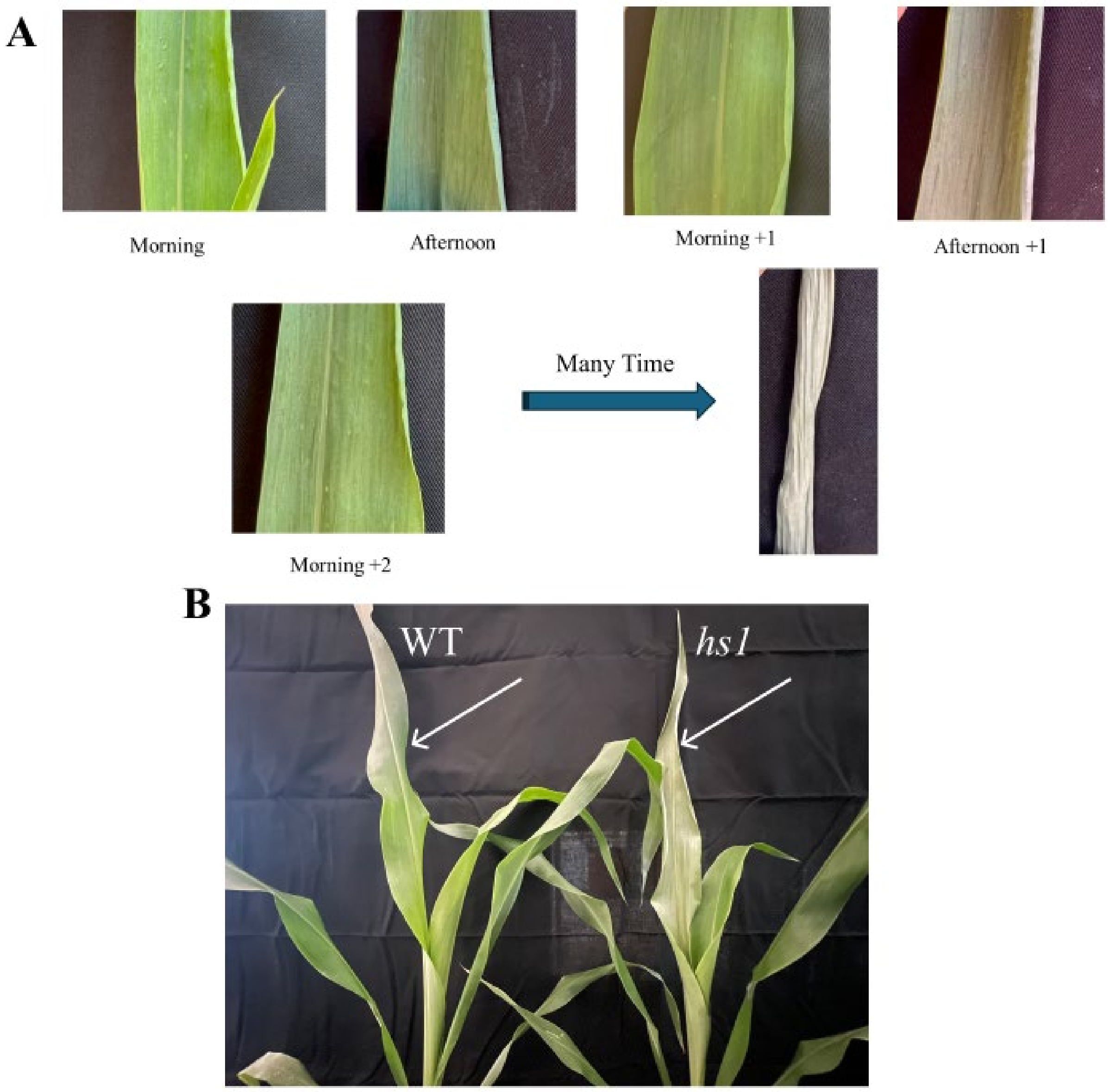


**Supplementary Figure S2.** Phenotypes of developing leaves in the field and greenhouse. **A)** Developing leaf of *hs1* during consecutive days in field condition (>34 °C). **B)** WT and *hs1* in the greenhouse under optimal conditions. White arrows indicate developing leaves. Optimal conditions: photosynthetic photon flux density (PPFD) 430–1162 μmol $m^{-2}$ $s^{-1}$; 26–30 °C.

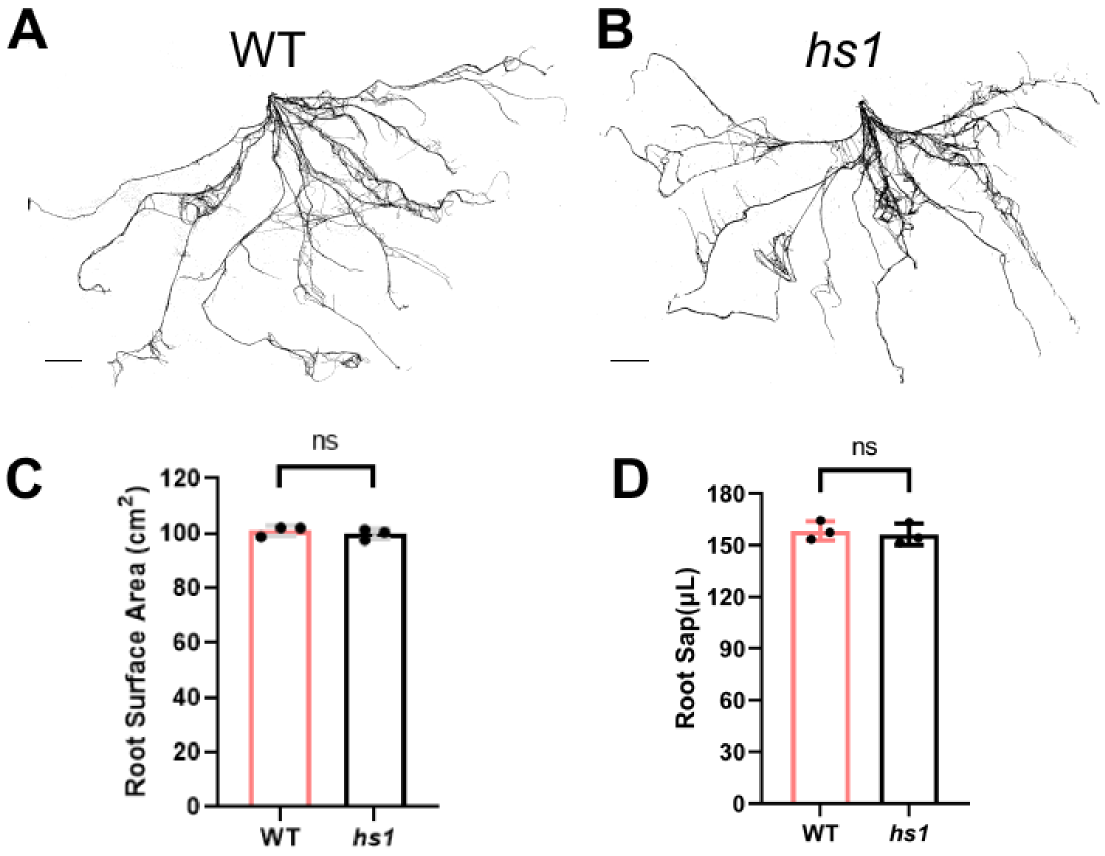


**Supplementary Figure S3.** Root traits of WT and *hs1* plants. **A, B)** Root morphology in WT (A) and *hs1* (B). Bar = 5cm. **C)** Quantification of root surface area in WT and *hs1*. **D)** Quantification of root sap (defined as root xylem sap exudate) in WT and *hs1*. For all statistical analyses, values are shown as the mean ± standard deviation from three biological replicates. ns, not significant ($P > 0.05$); two-tailed Student's t-test.

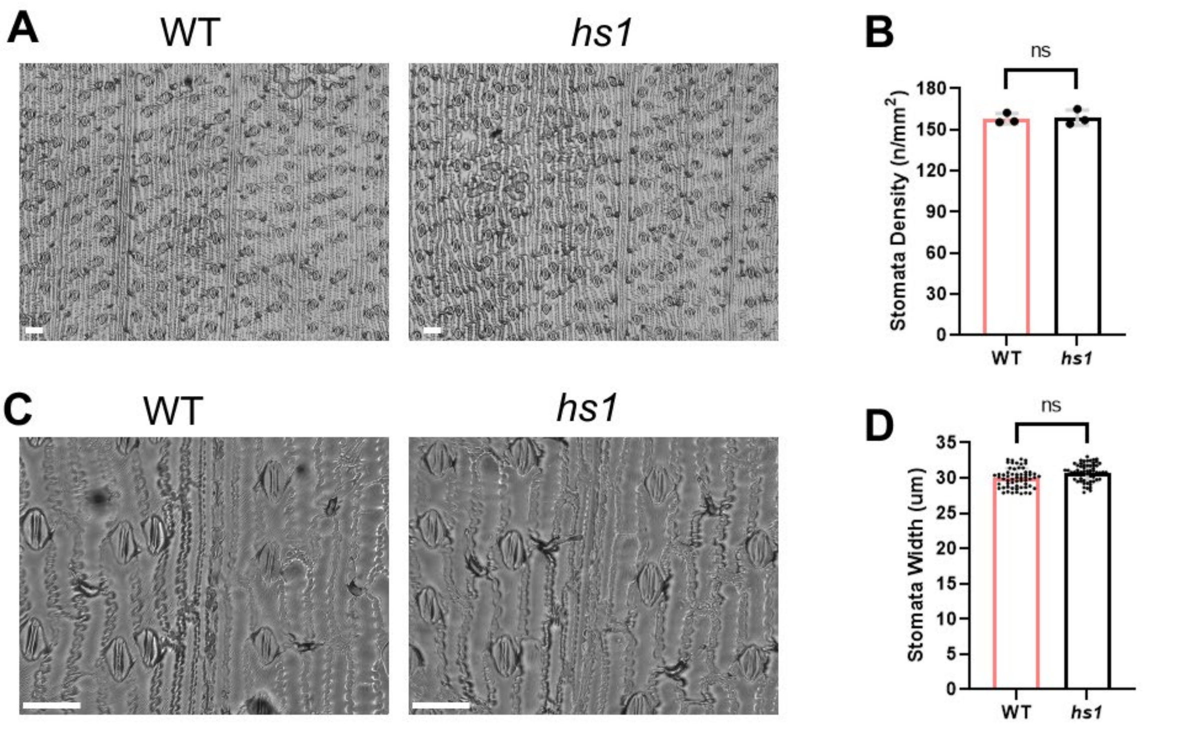


**Supplementary Figure S4.** Stomata traits of WT and *hs1* plants. **A)** Stomatal density in WT and *hs1* observed under a 100× light microscope. Bar = 50 μm. **B)** Quantification of stomatal density in WT and *hs1*. Values represent mean ± SD from three biological replicates. **C)** Stomatal morphology in WT and *hs1* observed under a 100× light microscope. Bar = 50 μm. **D)** Quantification of stomatal size in WT and *hs1*. Values represent mean ± SD from three biological replicates. For all statistical analyses: ns, not significant ($P > 0.05$); two-tailed Student's t-test.

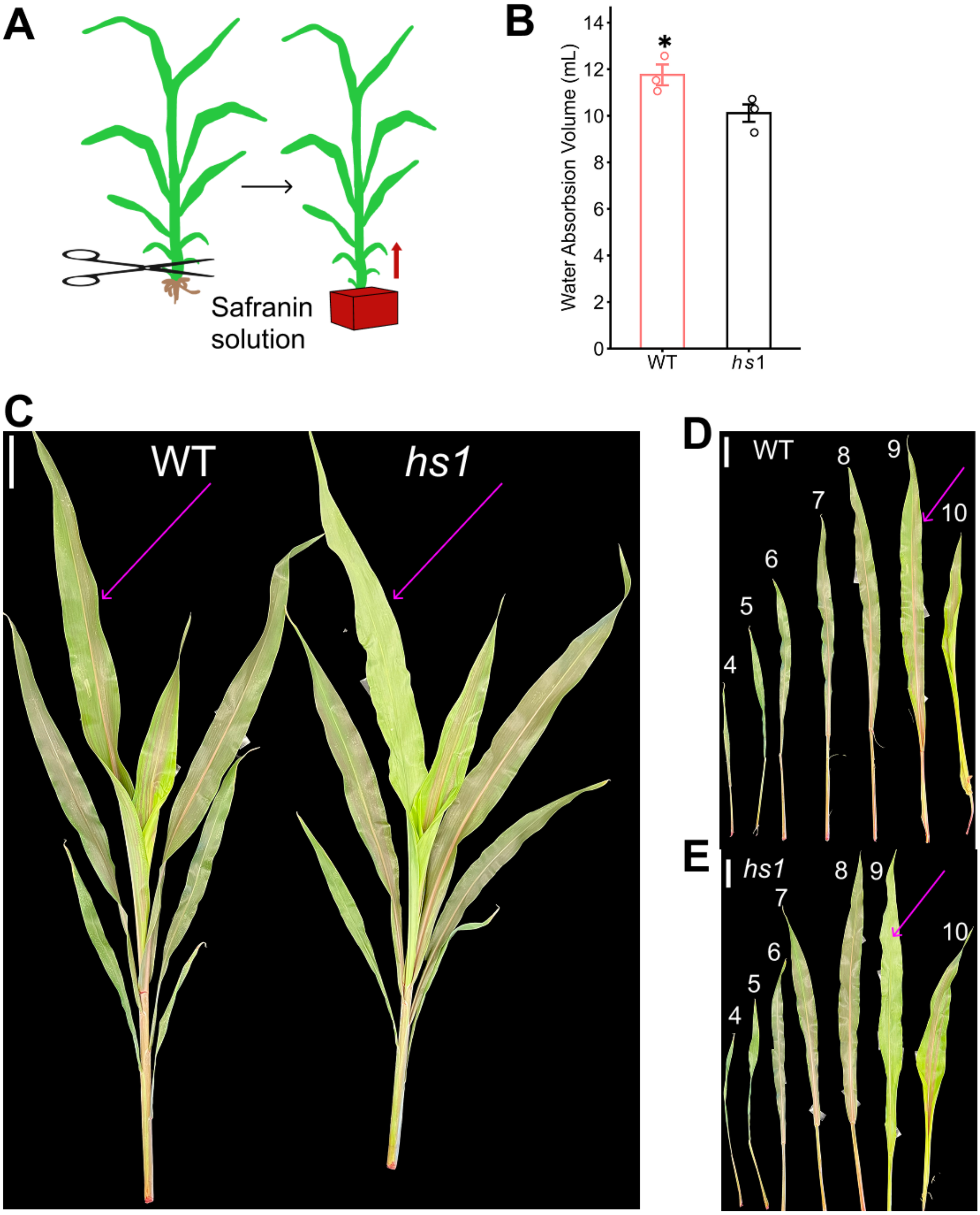


**Supplementary Figure S5.** Comparison of safranin dye transport in aerial tissues of WT and *hs1*. **A)** Schematic of the safranin uptake assay used to assess xylem transport. **B)** Statistical comparisons of safranin solution absorbed by WT and *hs1*. Values represent mean ± SD from three biological replicates. *P < 0.05 (two-tailed Student's t-test). **C)** Overview of safranin dye distribution in WT and *hs1* after 2 h. **D, E)** Safranin dye distribution in the $4^{th}$ to $10^{th}$ leaves of WT (D) and *hs1* (E) collected 2 h after assay. Arrows indicate developing leaves. Bar = 5 cm.

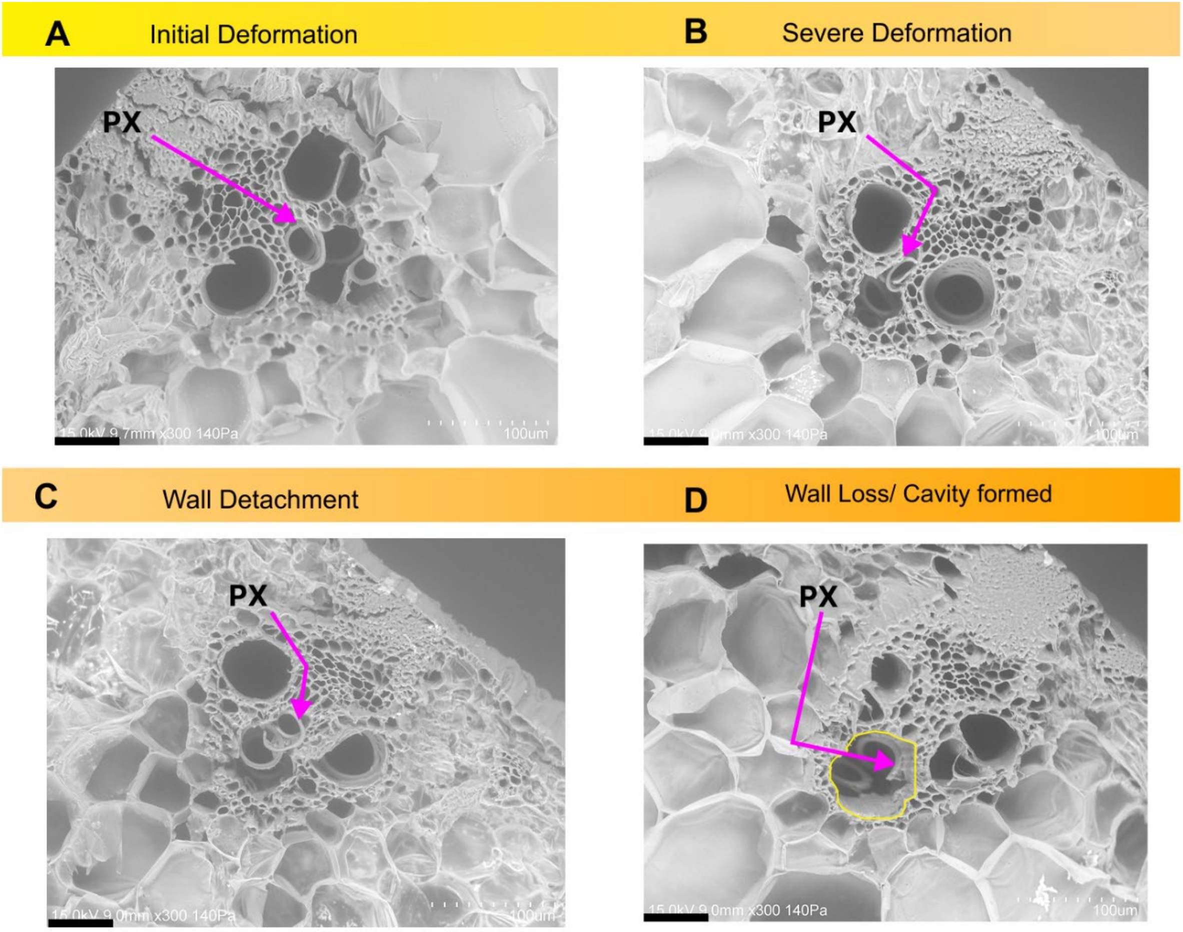


**Supplementary Figure S6.** Different collapse morphologies of protoxylem vessels (PX) in the midrib of developing leaf base of *hs1* observed by scanning electron microscopy (SEM). **A)** Initial mild deformation of PX in *hs1*. **B)** Severe deformation of PX in *hs1*. **C)** Secondary cell wall (SCW) extrusion in *hs1*. **D)** Cavity formation accompanied by SCW detachment and accumulation within the cavity in *hs1*. Arrows indicate representative PX; circles indicate cavities; bar = 500 µm.

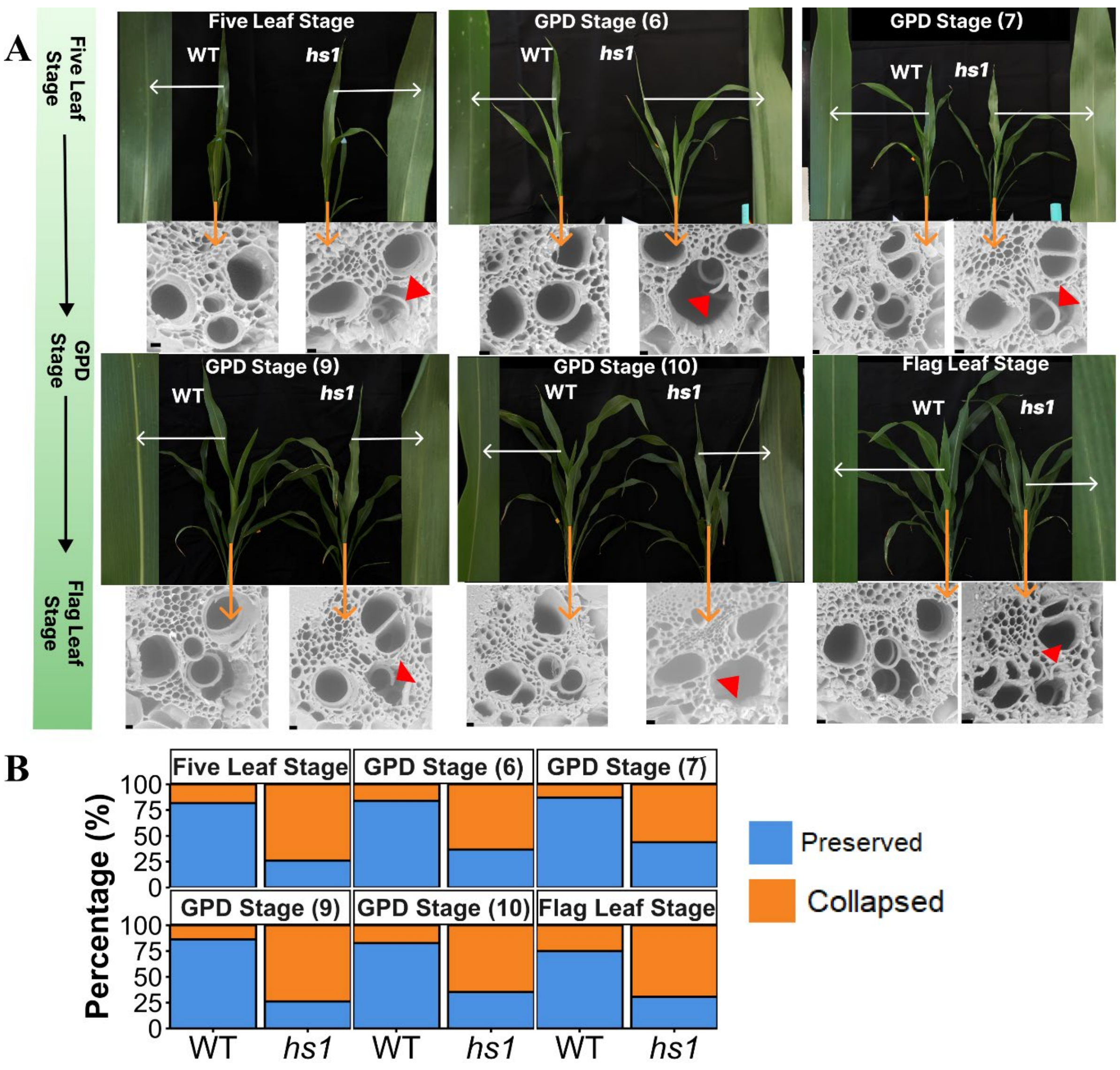


**Supplementary Figure S7**. Persistent protoxylem collapse in developing leaves across stages. **A)** Comparisons of WT and *hs1* from the five-leaf to the flag-leaf stage in the greenhouse. Enlarged views of developing leaves are shown at the side of each stage, with scanning electron microscopy (SEM) images of protoxylem (PX) in the corresponding leaf base midribs shown below. White arrows indicate developing leaves, orange arrows indicate corresponding vascular bundles there, and red triangles indicate collapsed PX. Bars = 10 µm. **B)** Quantification of PX integrity shown in (A). Data were obtained from three biological replicates per genotype. Individual PX vessels were pooled for analysis at each developmental stage (n = 13-44 PX vessels per genotype per stage). In (A) and (B), GPD (n), n denotes the number of expanded leaves.

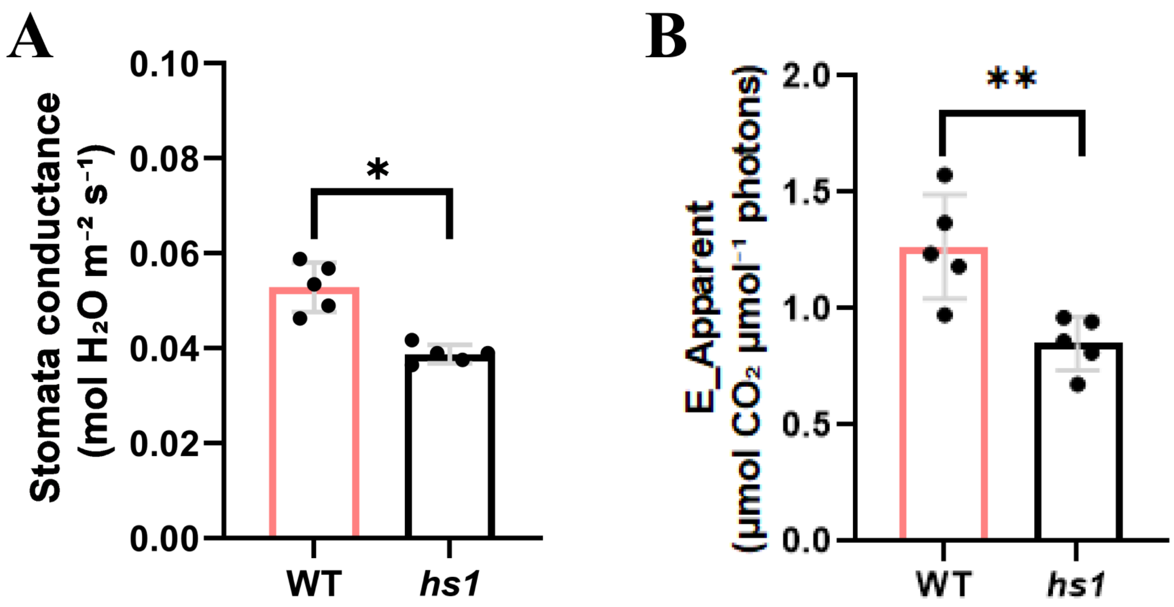


**Supplementary Figure S8**. Comparison of leaf transpiration parameters in developing leaves of WT and *hs1* in the morning. **A, B)** Comparison of stomatal conductance (A) and apparent transpiration rate (B) in developing leaves of WT and *hs1* in the morning. Values represent mean ± SD from five biological replicates; **P < 0.01, *P < 0.05 (two-tailed Student's t-test).

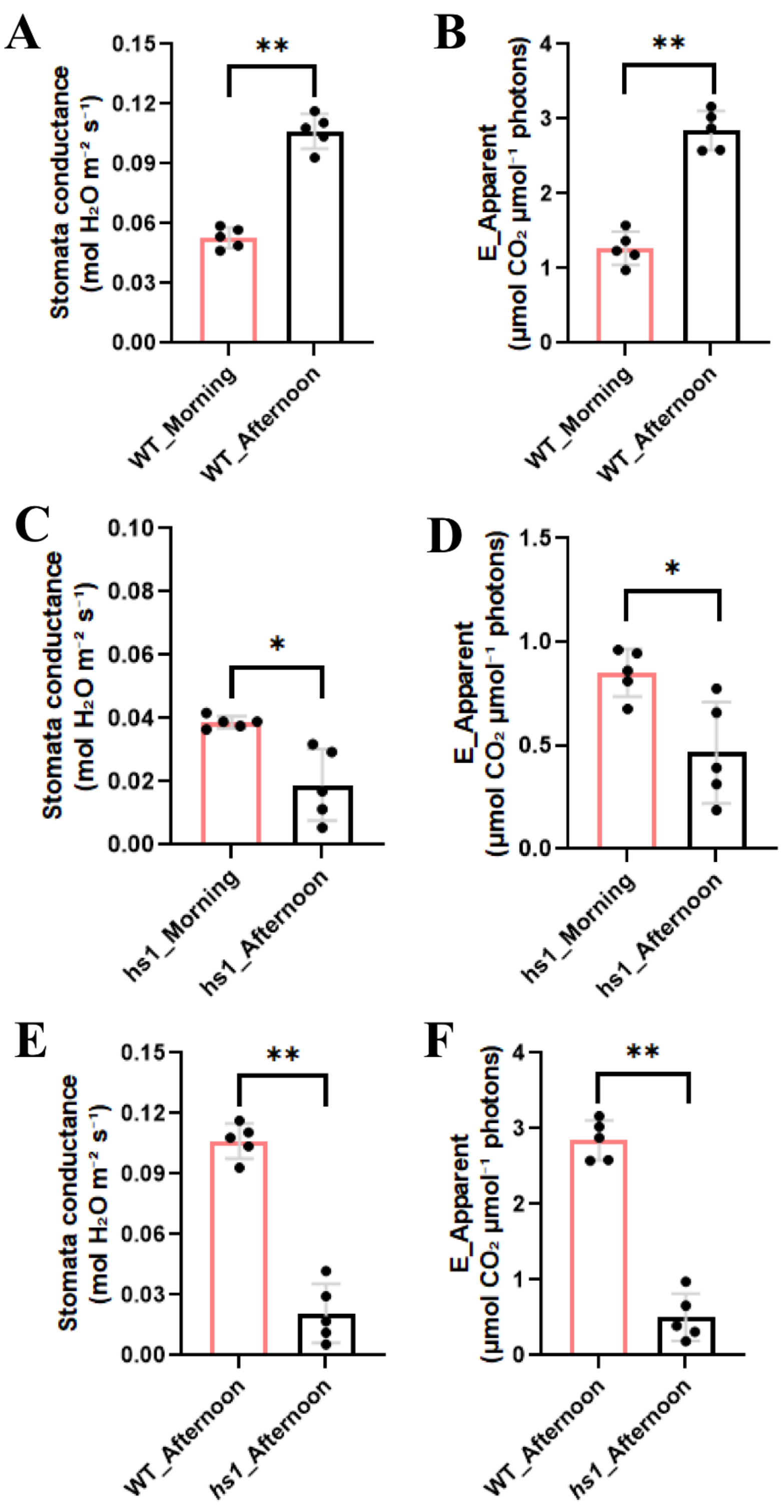


**Supplementary Figure S9**. Diurnal changes in leaf transpiration parameters in developing leaves of WT and *hs1*. **A to D)** Statistical comparisons of morning and afternoon stomatal conductance ($g_s$) and apparent transpiration rate (E) within WT (A, B) and *hs1* (C, D) developing leaves. **E, F)** Statistical comparisons of $g_s$ (E) and E (F) between WT and *hs1* plants in the afternoon. For all statistical analyses, values are shown as the mean ± standard deviation from five biological replicates. $^{**}P < 0.01$, $^{*}P < 0.05$ (two-tailed Student's t-test).

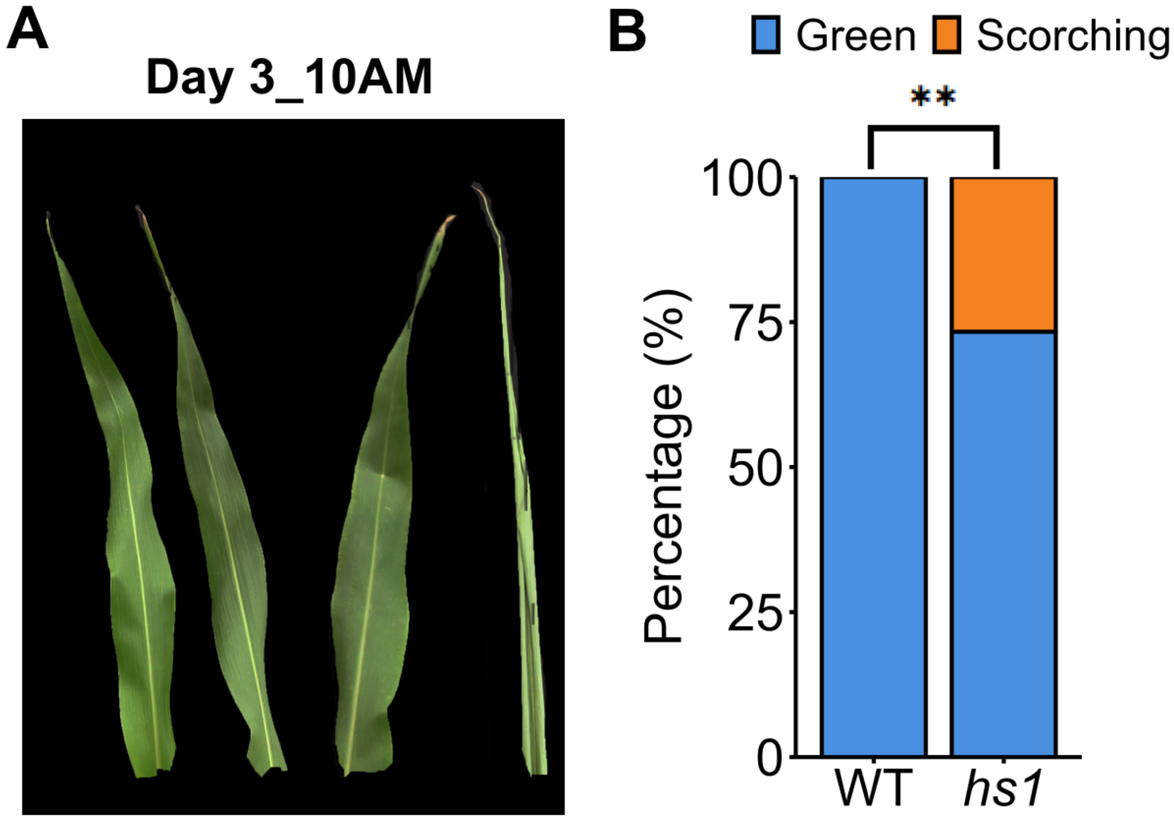


**Supplementary Figure S10**. Recovery rate of developing leaves in WT and *hs1* plants after two successive days. **A, B)** Developing leaves morphology (A) and recovery rate (B) of WT and *hs1* plants at 10AM after two days. Recovery rate was evaluated from 30 plants per genotype based on greenness of the leaves. Statistical significance was determined using Fisher's exact test (**P < 0.01).

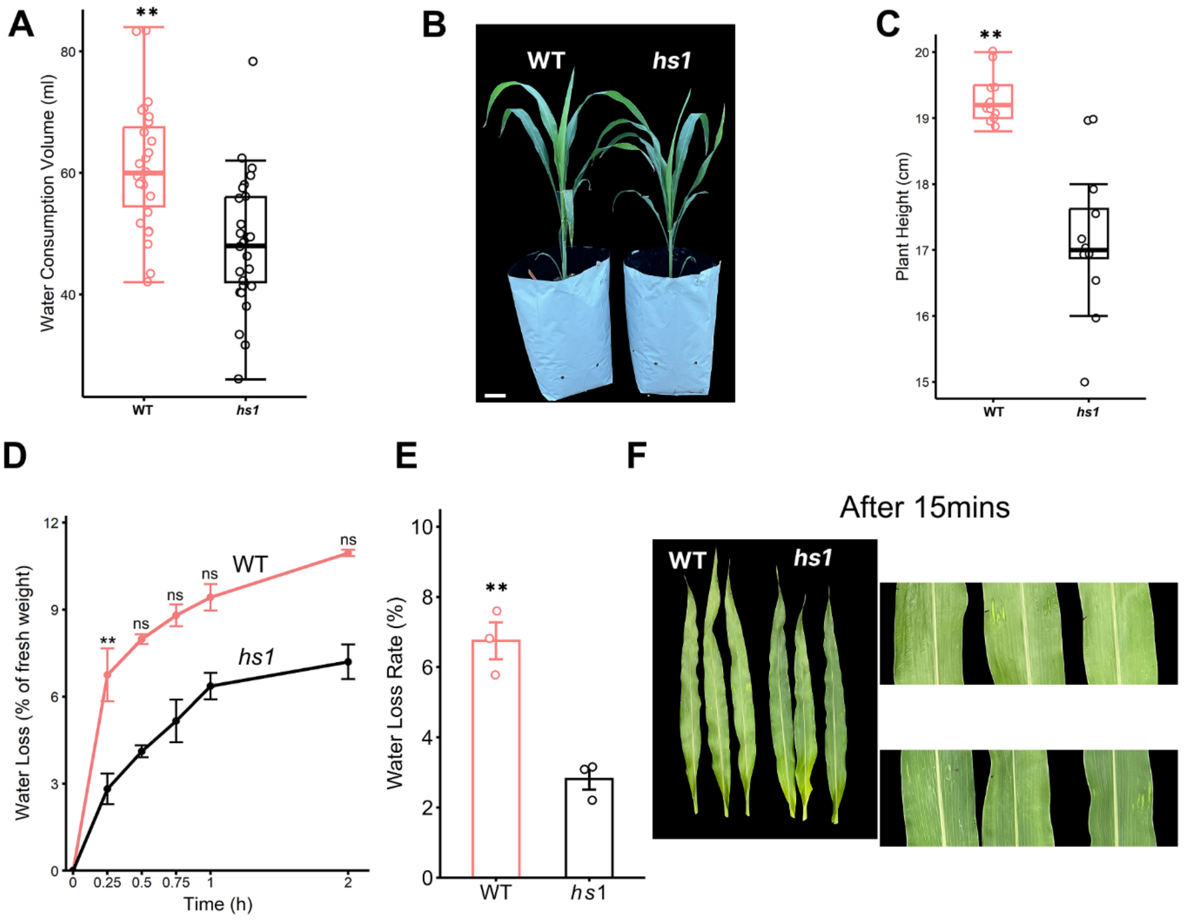


**Supplementary Figure S11**. Altered water consumption and dehydration responses in *hs1*. **A)** Statistical comparison of water consumption volume in WT and *hs1* over a 24 h period; boxes represent interquartile range; horizontal lines denote median; N = 26. **B)** Phenotypic comparison of plant height in WT and *hs1*; bar = 5 cm. **C)** Statistical comparison of plant height in WT and *hs1*; boxes represent interquartile range; horizontal lines denote median; N = 12. **D)** Statistical comparison of water loss over time from detached developing leaves of WT and *hs1*; points represent mean; error bars indicate SD; N = 3. **E)** Statistical comparison of water loss rate 15 min after detachment in WT and *hs1*; values represent mean ± SD; N = 3. **F)** Phenotypic comparison of detached leaves 15 min after detachment in WT and *hs1*; right panel shows an enlarged view of the central region of the left panel. For all statistical analyses: **P < 0.01; ns, not significant (P > 0.05); two-tailed Student's t-test.

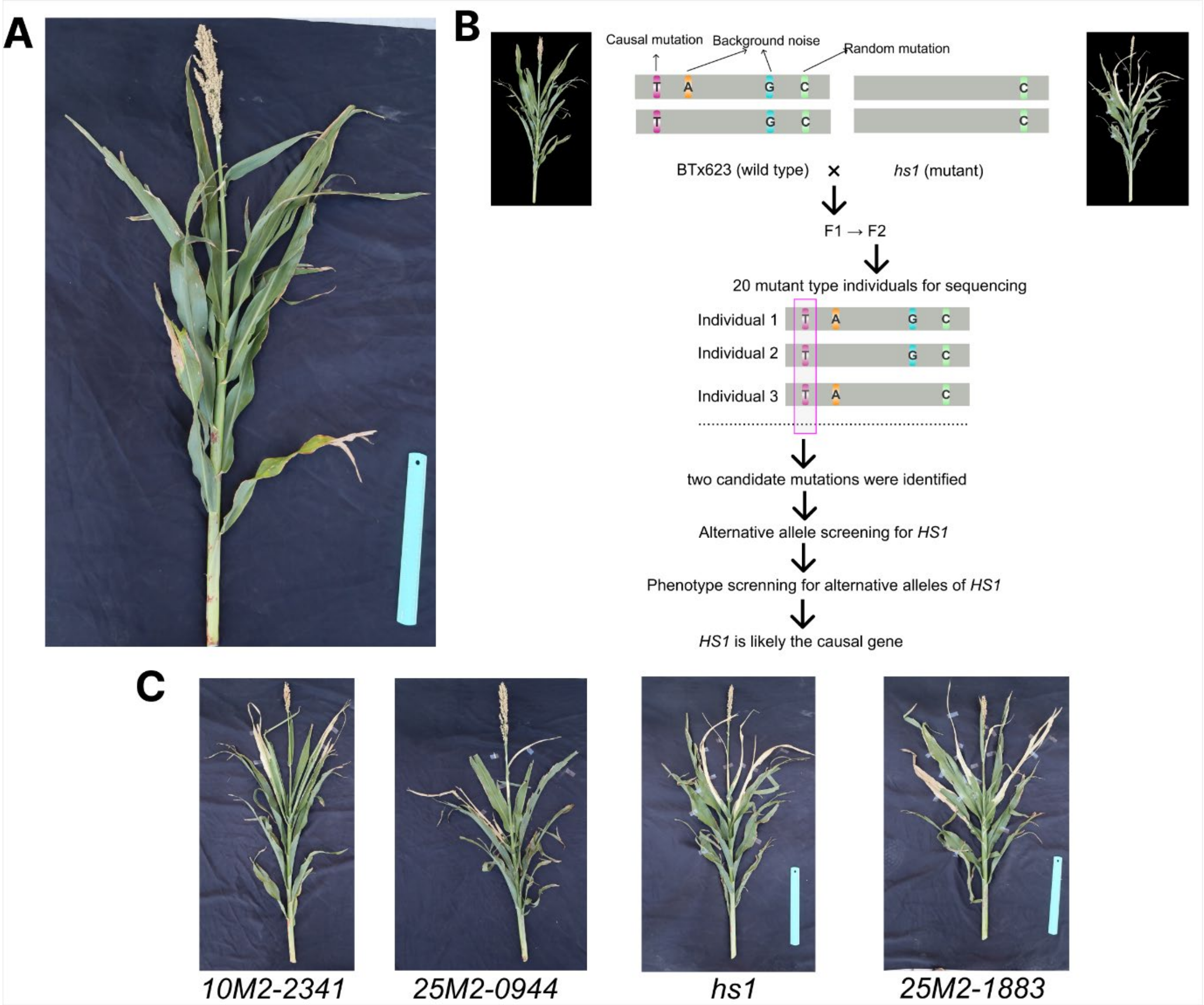


**Supplementary Figure S12.** Genetic mapping and allelic validation of *HS1*. **A)** Phenotype representative F1 plant derived from a cross between WT and *hs1* grown in the field. **B)** Workflow of bulk segregant analysis combined with whole-genome sequencing used to identify the causal mutation in the *hs1* F2 population. **C)** Phenotypes of four independent *HS1* alternative alleles (*10M2-2341*, *hs1*, *25M2-0944*, and *25M2-1883*).

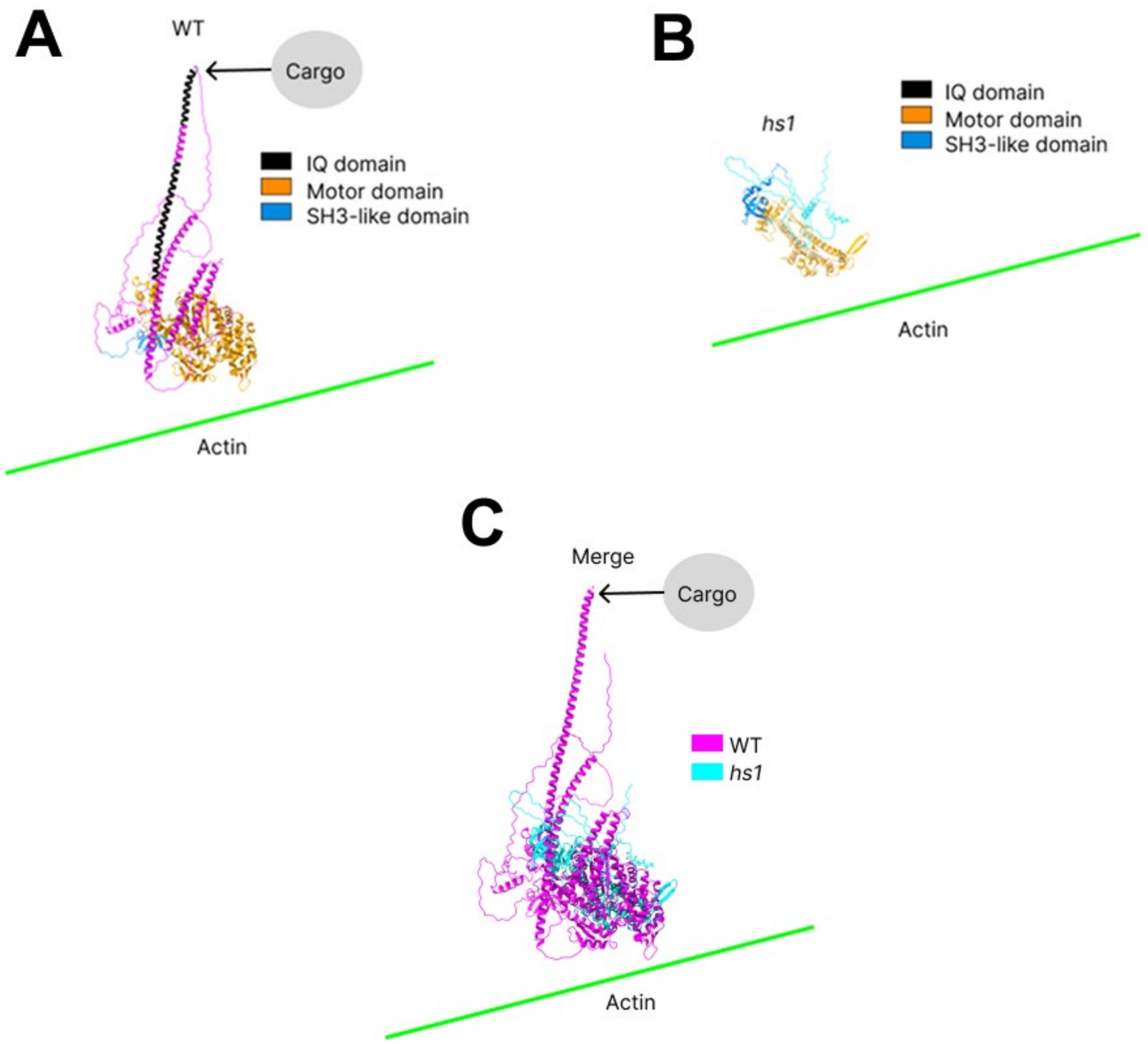


**Supplementary Figure S13.** Predicted structural alterations of HS1 in *hs1* relative to WT. **A)** Predicted three-dimensional structure of HS1 in WT. **B)** Predicted three-dimensional structure of HS1 in *hs1*. **C)** Overlay of predicted HS1 structures in WT and *hs1*.

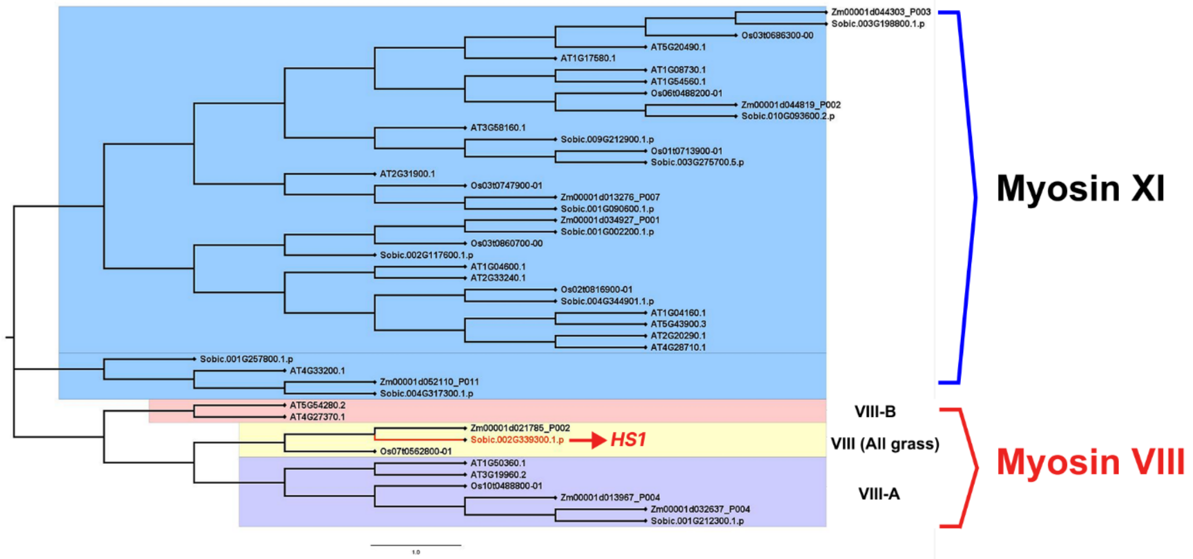


**Supplementary Figure S14.** Phylogenetic tree of myosin proteins in *Sorghum bicolor*, *Zea mays*, *Oryza sativa*, and *Arabidopsis thaliana.*

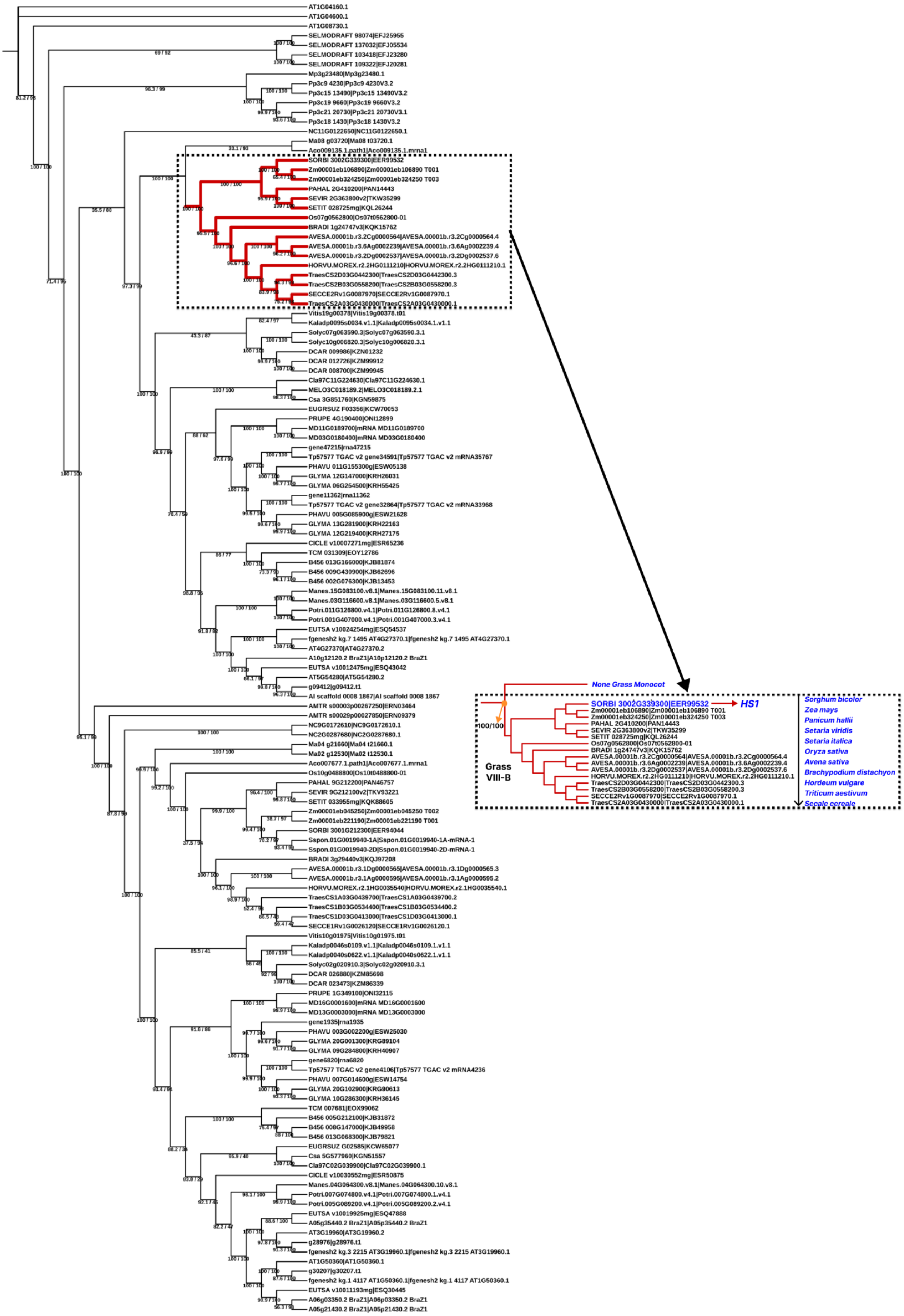
None Grass Monocot
SORBI 3002G339300|EER99532
HS1
100/100
Grass
VIII-B
Sorghum bicolor
Zea mays
Panicum hallii
Setaria viridis
Setaria italica
Oryza sativa
Avena sativa
Brachypodium distachyon
Hordeum vulgare
Triticum aestivum
Secale cereale

**Supplementary Figure S15.** Maximum-likelihood (ML) phylogeny of the myosin gene family. SH-aLRT/bootstrap support values are shown at the nodes. The grass-specific VIII-B clade is highlighted in red. The right panel presents a subtree of plant myosin VIII proteins, with *HS1* from Sorghum bicolor indicated

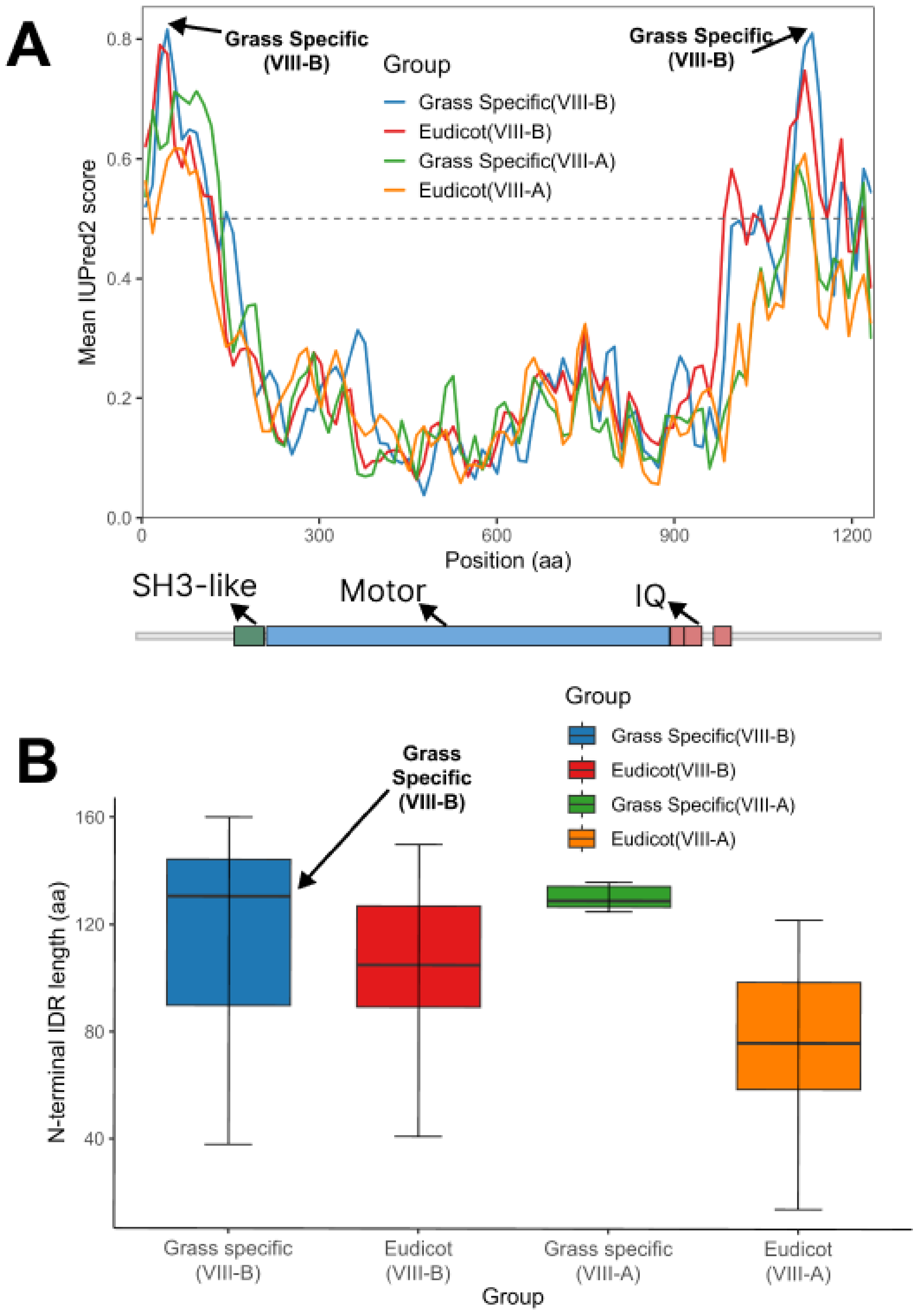


**Supplementary Figure S16.** Sequence features of the grass-specific VIII-B clade at terminal regions. **A)** Comparative intrinsic disorder (IDR) profiles of myosin VIII proteins across major plant lineages. The schematic below shows conserved domain architecture aligned to the IDR profiles above, using *HS1* as the reference protein. **B)** Comparison of N-terminal IDR length across major plant lineages.

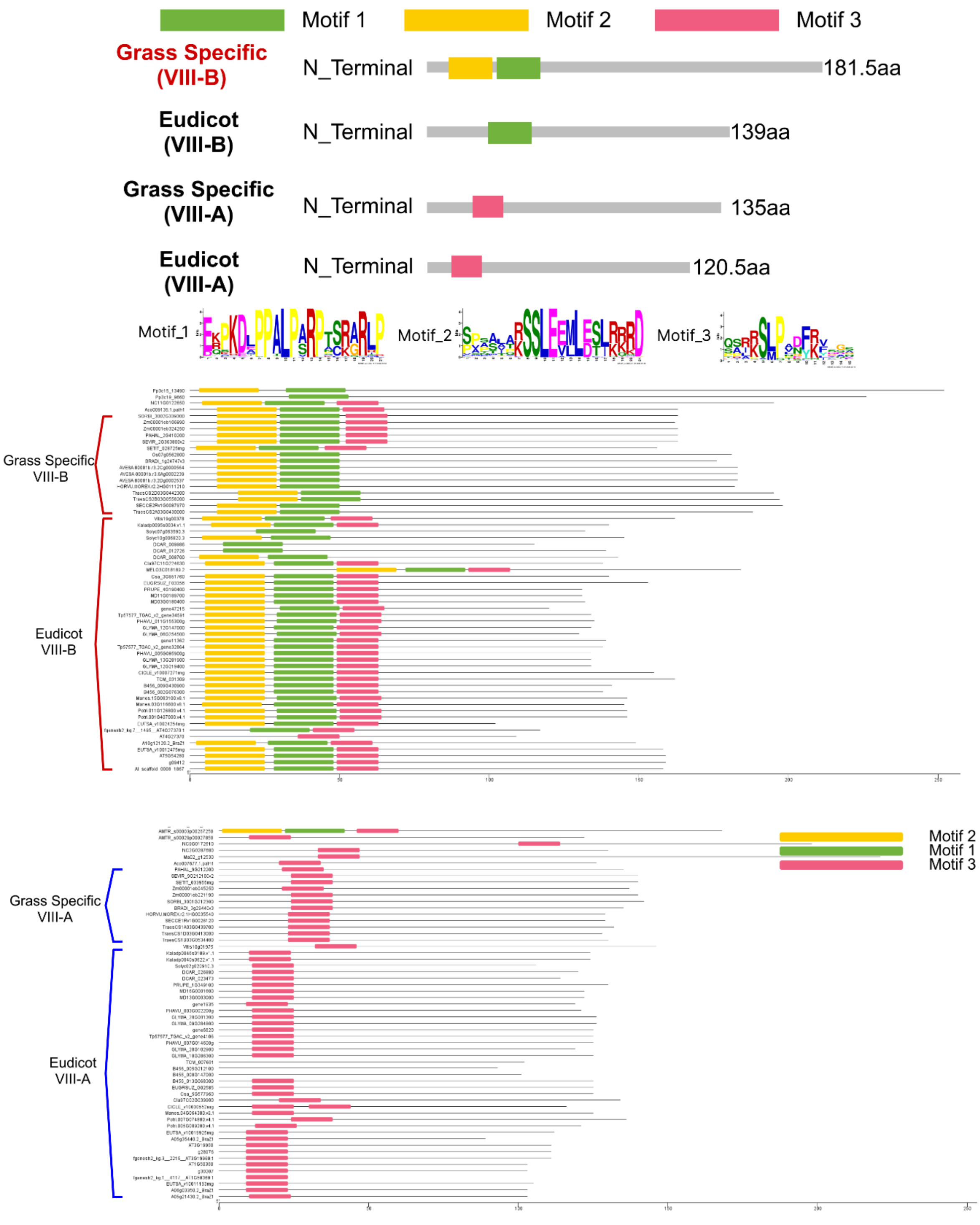


**Supplementary Figure S17**. Features of myosin VIII protein N-terminal sequences. **A)** Summary schematic of N-terminal features, illustrating the overall motif organization shown in B and C; conserved motifs are defined as present in >90% of members within the corresponding clade; sequence logos of Motif 1–3 are shown below. **B)** N-terminal motif distribution and length in myosin VIII-B proteins across major plant lineages. **C)** N-terminal motif distribution and length in myosin VIII-A proteins across major plant lineages.

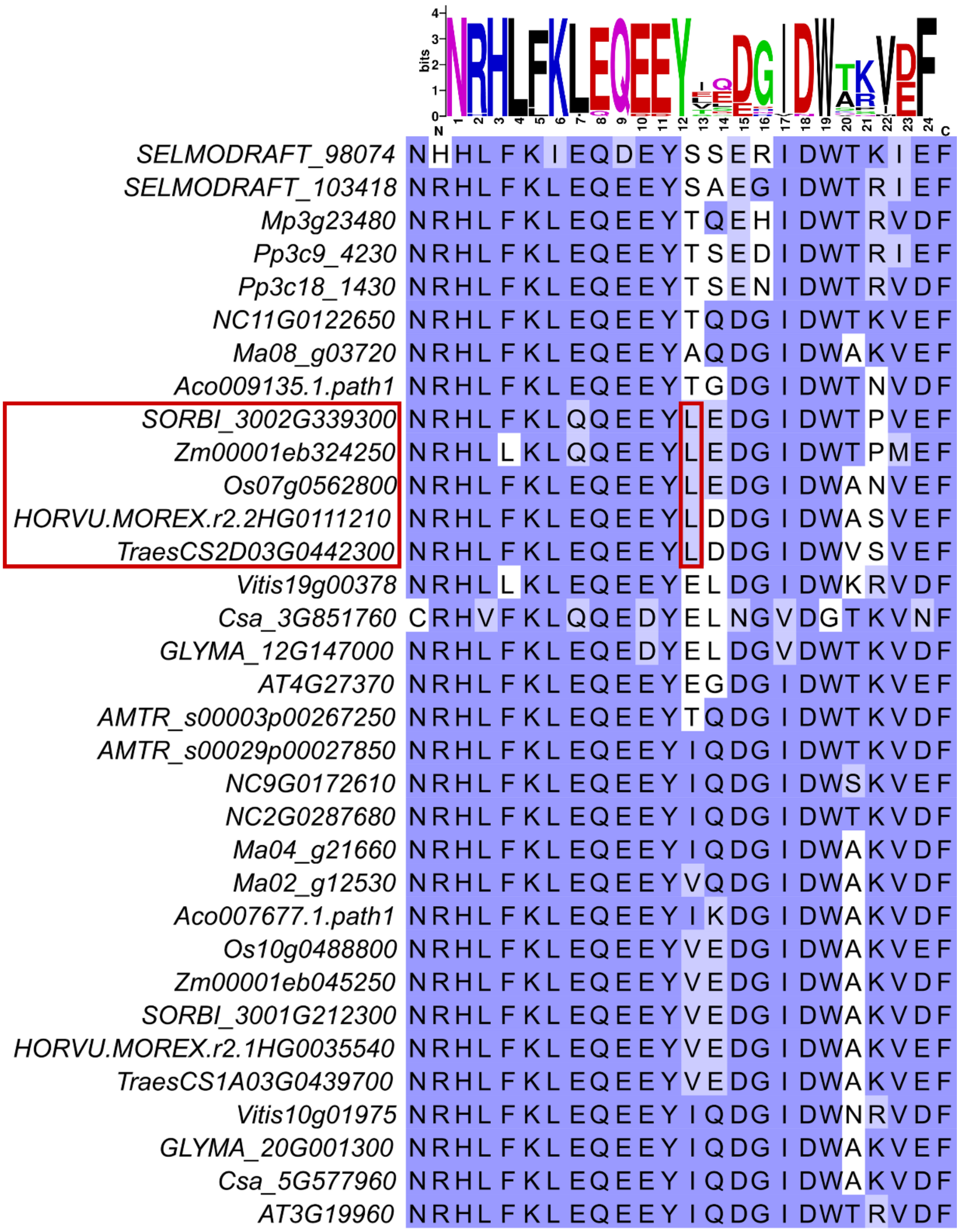


**Supplementary Figure S18.** Multiple sequence alignment of the relay loop region. Red boxes indicate the grass-specific VIII-B clade and the uniquely conserved leucine (L).

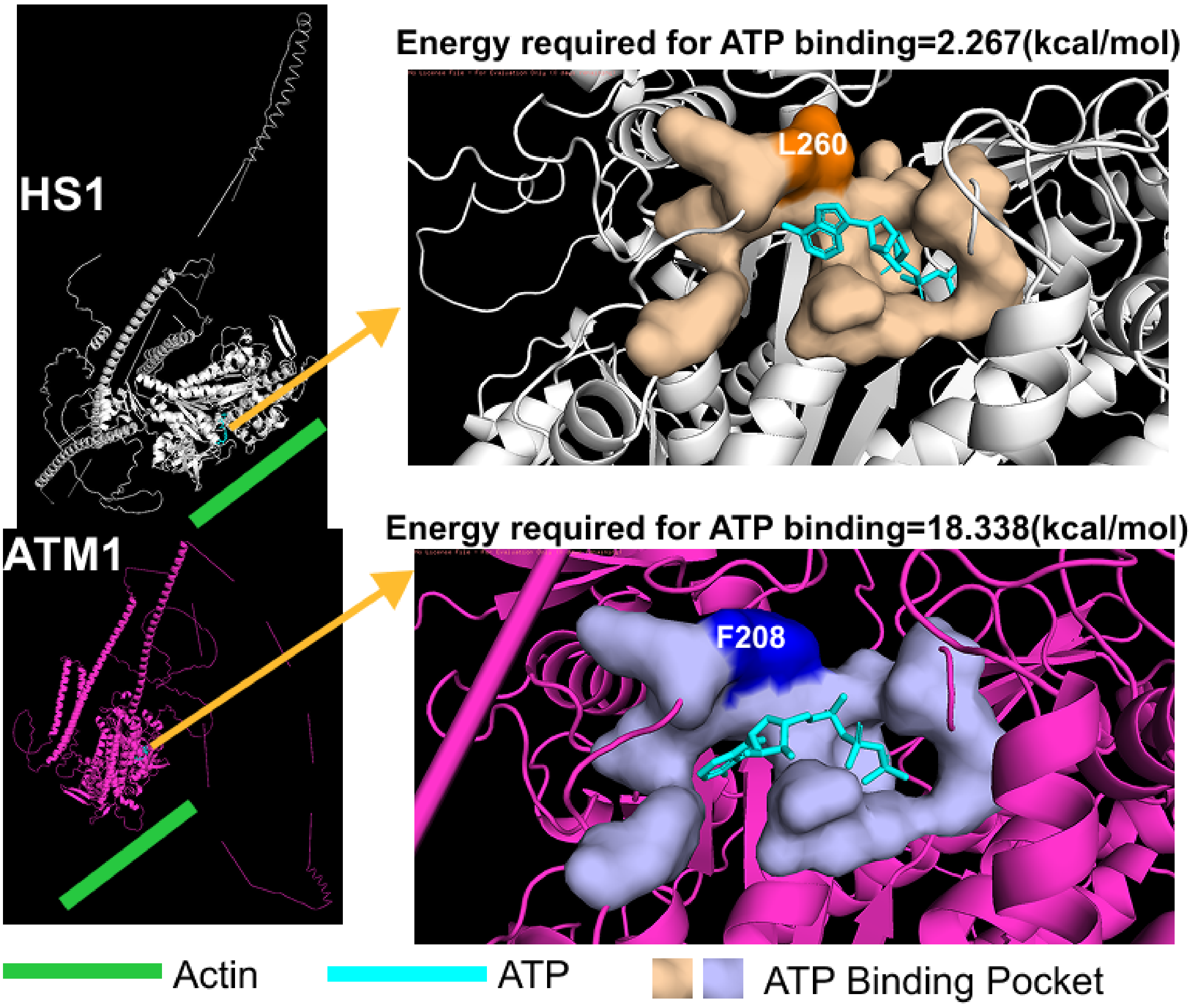


**Soybean myosin VIII-B: Energy required for ATP binding = 6.677 kcal/mol**

**Supplementary Figure S19.** Structural comparison of predicted ATP-binding properties of grass VIII-B and eudicot VIII-A proteins. *HS1* is shown as a representative of grass VIII-B, and *Arabidopsis thaliana* MYOSIN 1 (*ATM1*) as a representative of eudicot VIII-A. ATP (cyan), actin (green), and the ATP-binding cavity surface are displayed. Residue L260 in *HS1* and residue F208 in *ATM1* are indicated.

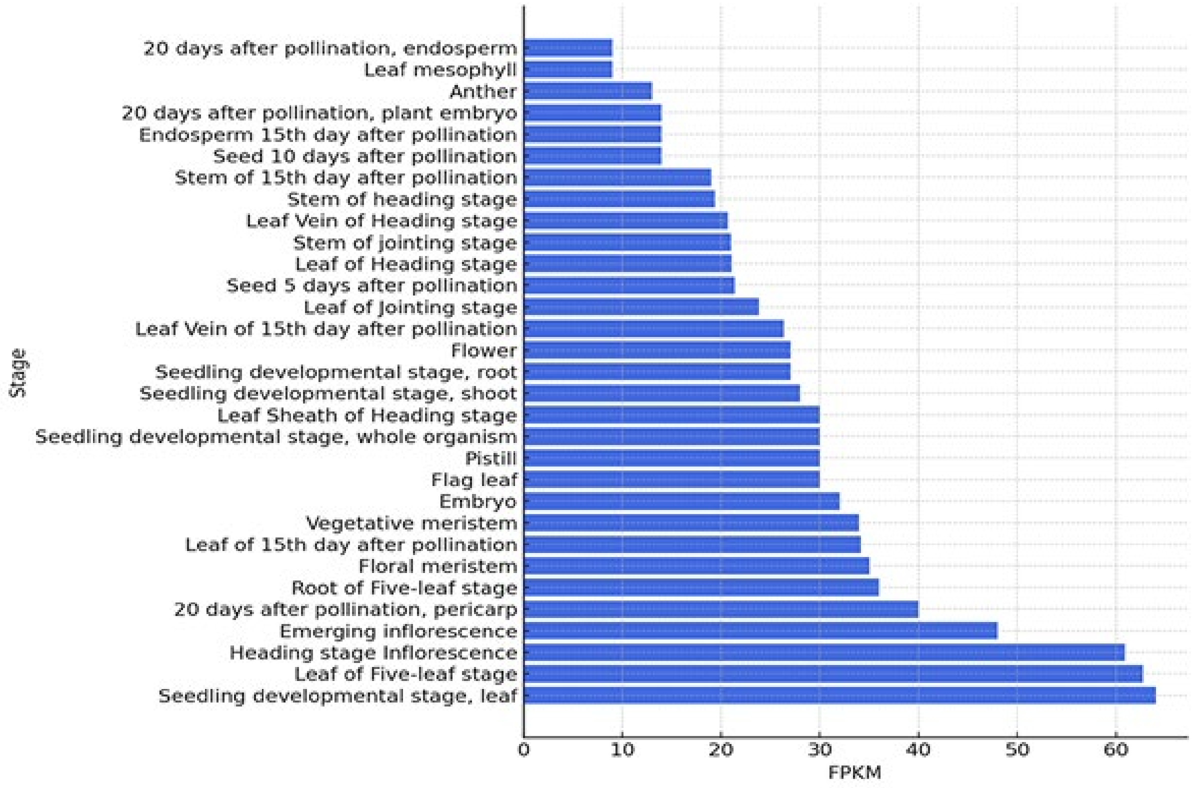


**Supplementary Figure S20.** Transcript levels FPKM (Fragments Per Kilobase of transcript per Million mapped reads) of *HS1* in different sorghum tissues. Data was extracted from public databases of Ensembl Expression Atlas (Papatheodorou et al., 2018) and Phytozome (Goodstein et al., 2012).

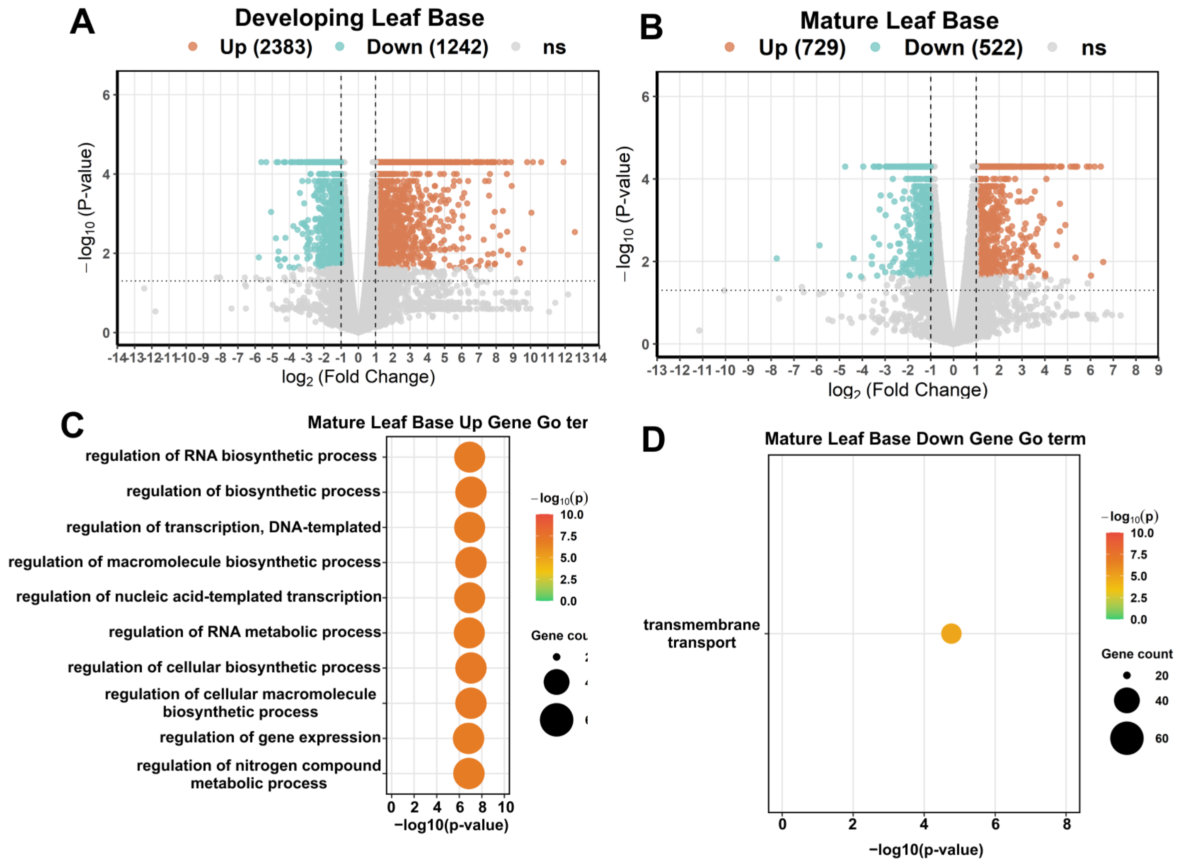


**Supplementary Figure S21.** Transcriptome analysis of the leaf base midrib in WT and *hs1*. **A, B)** Number of differentially expressed genes (DEGs) in immature leaves (A), and mature leaves (B) of *hs1* compared with WT. **C, D)** GO enrichment analysis of upregulated (C) and downregulated (D) genes in mature leaf base midrib of *hs1* compared with WT. The top 10 GO terms ($P < 0.01$), ranked by p-value.

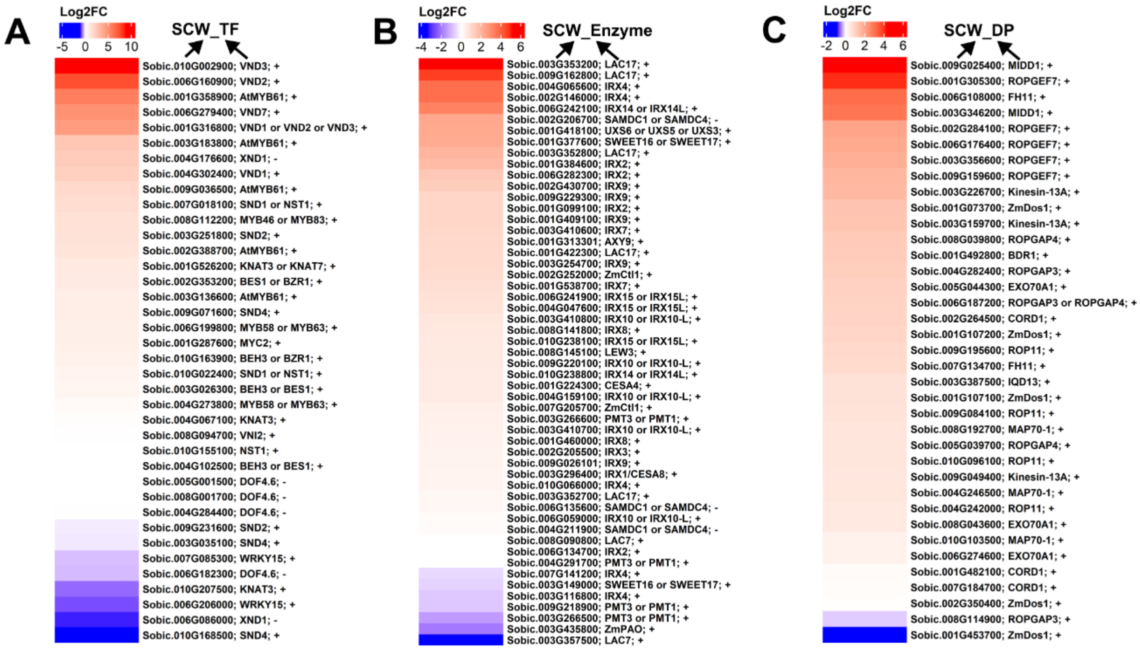


**Supplementary Figure S22.** Heatmaps showing $\log_2$ fold changes in sorghum ortholog expression in *hs1* relative to WT for known secondary cell wall (SCW)–related genes from *Arabidopsis thaliana*, *Zea mays*, and *Oryza sativa*, grouped as transcription factors (SCW_TF) (A), biosynthesis enzymes (SCW_Enzyme) (B), and deposition-related genes (SCW_DP) (C). This figure presents the complete dataset corresponding to Figs. 6, D to F, including all detected genes without |log2 fold change| filtering. "+" and "–" indicate positive and negative regulators of SCW formation, respectively; *q-value* < *0.05*.

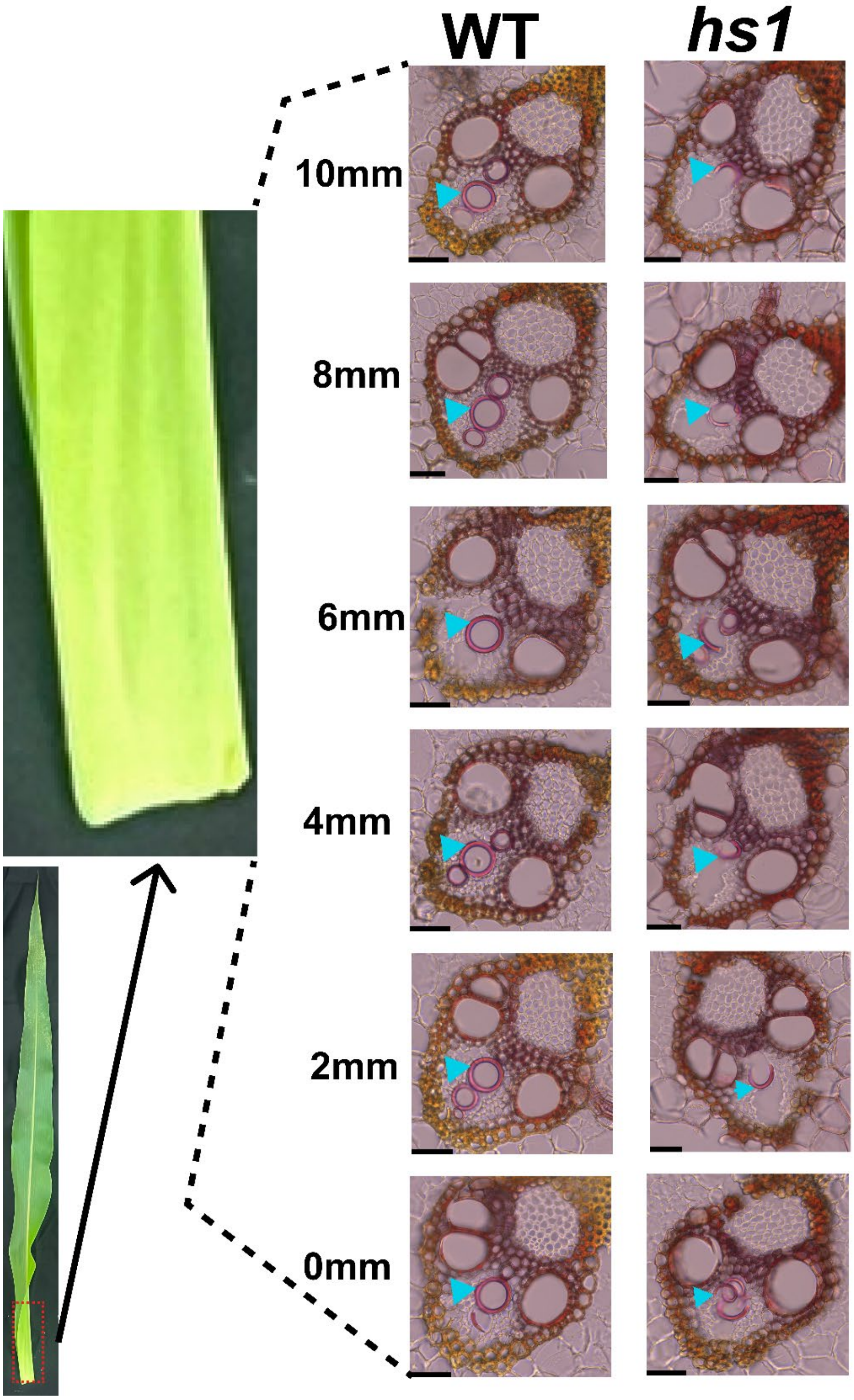


**Supplementary Figure S23**. Serial transverse sections of the developing leaf base midrib at indicated distances (0–10 mm) above the leaf sheath in WT and *hs1*. Protoxylem vessels are indicated by cyan triangles. Bar = 50 µm.

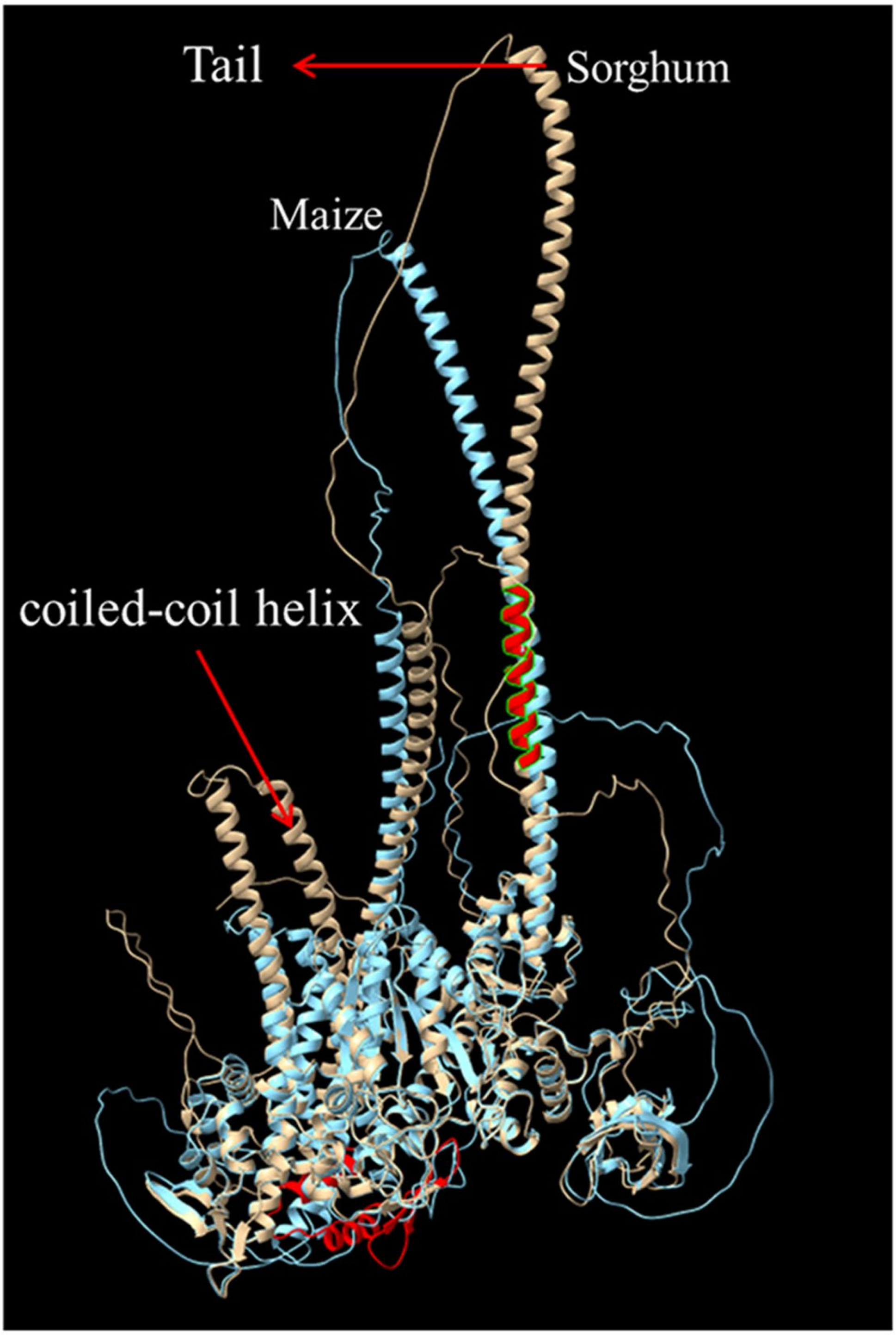


**Supplementary Figure S24**. Protein three-dimensional structure alignment of *HS1* and *ZmVIII-3*. Red circles indicate regions corresponding to multiple 15-aa gaps between *HS1* and *ZmVIII-3* in the protein sequence alignment.
